# S-adenosylmethionine metabolism buffering is regulated by glycine N-methyltransferase decrease via nuclear ubiquitin-proteasome system

**DOI:** 10.1101/2024.08.21.609067

**Authors:** Soshiro Kashio, Masayuki Miura

## Abstract

Metabolic homeostasis is essential for survival; however, many studies have focused on the fluctuations of these factors. Furthermore, while metabolic homeostasis depends on the balance between the production and consumption of metabolites, there have been limited investigations into the mechanisms regulating their consumption. S-adenosylmethionine (SAM) metabolism has diverse functions, including methylation, polyamine biosynthesis, and transsulfuration, making its regulation and control crucial. Recent studies have revealed the feedback regulation of SAM production; however, the mechanisms governing its consumption are still poorly understood.

In this study, we focused on the stability of SAM levels in the fat body (FB) of *Drosophila*, which serves as a functional equivalent of the mammalian liver and adipose tissue, under conditions of SAM shortage, including nutrient deprivation. We found that glycine N-methyltransferase (Gnmt), a major SAM-consuming methyltransferase in the FB, decreased via the nuclear ubiquitin-proteasome system (UPS), along with the inhibition of SAM synthesis and starvation. The inhibition of Gnmt degradation by suppression of the nuclear UPS causes starvation tolerance. Thus, the regulation of Gnmt levels through nuclear UPS-mediated degradation helps maintain SAM levels under SAM shortage conditions.

**Significance Statement:** S-adenosylmethionine (SAM) metabolism is crucial for diverse functions, which are mediated through methylation process. Although the feedback regulation of SAM production has been explored extensively, our understanding of the mechanism behind SAM consumption remains incomplete. Constant levels of SAM have been observed in *Drosophila* fat bodies even under conditions of SAM shortage, including nutrient deficiency and inhibition of SAM synthesis. SAM levels are controlled by the degradation of glycine N-methyltransferase (Gnmt), a cytosolic SAM-consuming enzyme, via the nuclear ubiquitin-proteasome system under conditions of SAM shortage. Additionally, the inhibition of Gnmt degradation by suppression of the nuclear UPS causes starvation tolerance. Considering that SAM accumulation promotes energy expenditure *in vivo*, the starvation-dependent mechanism of Gnmt degradation is important for energy homeostasis.

## Introduction

Metabolism is a defining element of life. Harmonized metabolic homeostasis is indispensable for sustaining life, and its regulatory mechanisms have been extensively studied across various biological contexts and species. While the importance of maintaining a balance between the production and consumption of metabolites for effective feedback control is recognized(1), most studies have focused on variations in the levels of these factors, but the constant factors are often overlooked. In addition, research on metabolic stability mechanisms is primarily concentrated on production control, with reports pertaining to consumption-related control being limited.

S-adenosylmethionine (SAM) metabolism is a crucial process with diverse functions, in methylation, polyamine biosynthesis, and transsulfuration (TSF)(2). As a methyl donor, SAM participates in metabolic pathways originating from the amino acid methionine (Met) and circulates within the methionine cycle. It is utilized by various methyltransferases for the methylation of DNA, RNA, histones, phospholipids, and other substrates. During methylation, SAM is converted to S-adenosylhomocysteine (SAH) and is involved in polyamine biosynthesis and the Met salvage pathway. Furthermore, in the TSF pathway, SAM contributes via its metabolites SAH and homocysteine (Hcy). Hcy is converted to cystathionine (Cysta), cysteine (Cys), and glutathione (GSH) to regulate redox processes. Importantly, the conversion of Hcy to Met affects the folate cycle.

Recent studies have elucidated the feedback mechanisms regulating SAM production (3, 4). In *Caenorhabditis elegans*, under high-nutrient conditions, adenine methylation at the 3’ splice site of SAM synthetase (SAMS-3) inhibits splicing, resulting in a decrease in SAM levels. Conversely, under low-SAM conditions, SAMS-3 splicing progresses, leading to SAM production from Met. Additionally, translation-level control of SAM has also been observed in yeasts. SAM synthetase (Sams) in yeast features a riboswitch structure in its 5’ untranslated region (5’UTR)(5). When SAM is depleted, N6-methyladenosine in the 3’ UTR of *MAT2A*, mammalian SAM synthetase is reduced, leading to the stabilization of *MAT2A* mRNA (6). These findings highlight the multifaceted regulation of SAM production. Considering natural conditions such as nutrient deficiency, further research is required to explore the regulation of SAM consumption.

In mammals, the liver is the central organ for SAM metabolism, whereas in invertebrates the fat body (FB), which corresponds to the liver and white adipose tissue, is the key regulatory tissue for SAM metabolism. Notably, the knockdown of *sams* in *Drosophila* larval FB did not affect SAM levels (7). This suggests the existence of a distinct mechanism controlling SAM consumption. Glycine N-methyltransferase (Gnmt), which is highly expressed in the liver in mammals and FB in *Drosophila*, is an important regulator of SAM levels. Using SAM as a methyl donor, Gnmt converts glycine to sarcosine (Sar), which is subsequently converted back to glycine by sarcosine dehydrogenase. Gnmt protein levels in the FB unexpectedly decreased under *sams-RNAi* (7), suggesting that a decrease in Gnmt may help maintain SAM levels when SAM production is reduced.

In this study, we explored the mechanism of Gnmt downregulation due to SAM deficiency and identified the involvement of a specific ubiquitin ligases located in the nucleus. This regulatory mechanism of Gnmt degradation is important during starvation.

## Results

### Stability of SAM levels during starvation in larvae

First, we examined the metabolic state changes during nutrient deprivation by measuring water-soluble metabolites in *Drosophila* larvae under starvation conditions (Fig. 1A). Xanthine, uric acid, and kynurenine (Kyn) were among the metabolites that increased, whereas the levels of threonine (Thr), aspartic acid (Asp), and asparagine (Asn) decreased (Fig. 1B). Notably, the levels of metabolites associated with Met metabolism, such as SAM and 5’-methylthioadenosine (MTA), did not change even under starvation conditions (Fig. 1B and C and Table S1). In particular, examination of Met metabolism under starvation showed decreases in Met, SAH, and Cysta, whereas the levels of methionine sulfoxide increased, and the levels of SAM and MTA remained unchanged (Fig. 1C and D). These results suggest that SAM levels are maintained even under starvation conditions and likely contribute to the production of downstream metabolites and methylation processes.

**Figure 1.**
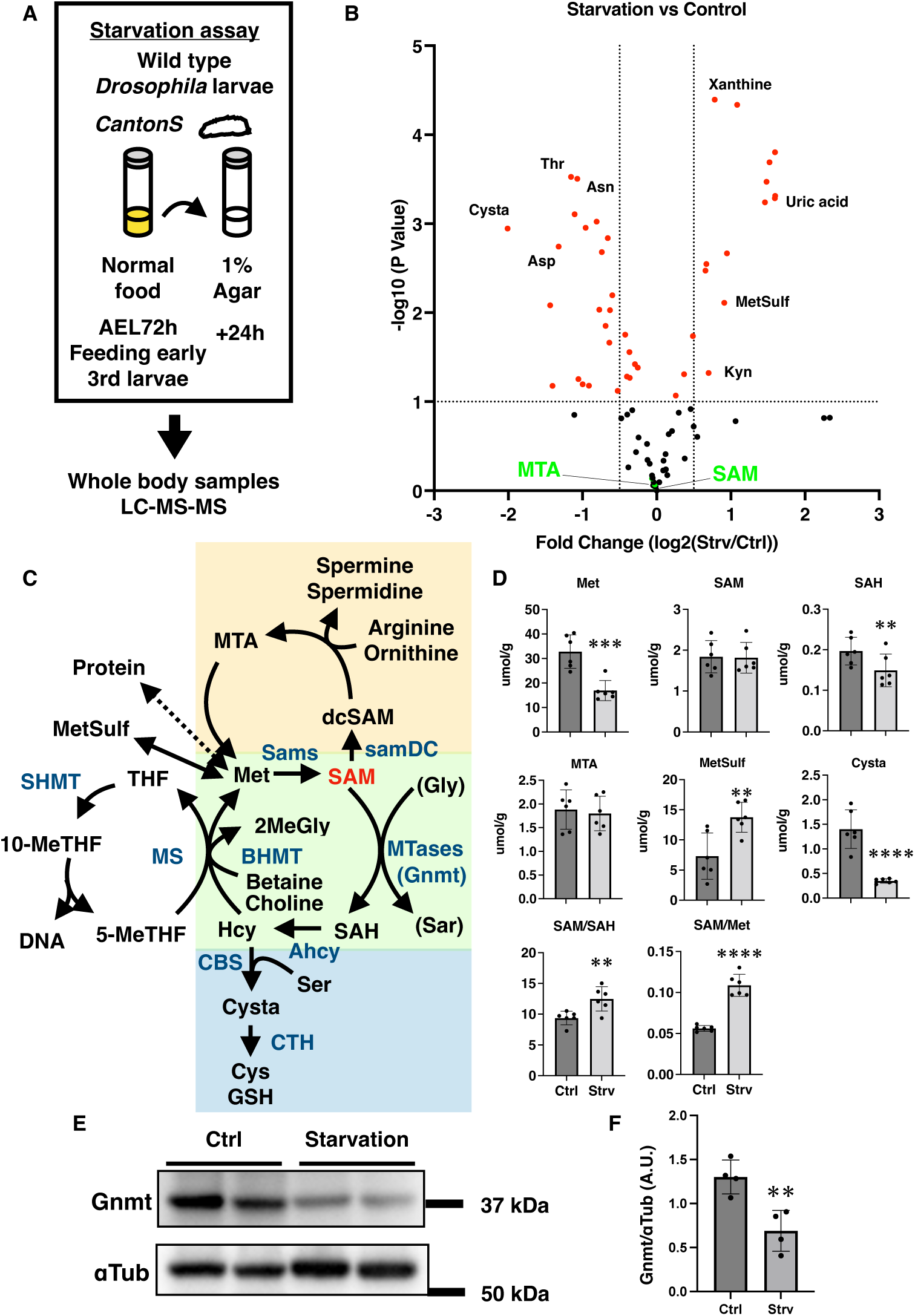
Stability of SAM levels during starvation in *Drosophila* larvae. (**A**) Schematic of the starvation assay for larvae and metabolic analyses. (**B**) Changes in metabolite levels during starvation. (**C**) Schematic of the SAM metabolism map. SAM: S-adenosylmethionine, SAH: S-adenosyl homocysteine, Hcy: homocysteine, Met: methionine, Cysta: cystathionine, Cys: cysteine, GSH: glutathione, Gly: glycine, Sar: sarcosine, MTA: 5′-methylthioadenosine, dcSAM: decarboxylated SAM, 2MeGly: dimethylglycine, THF: tetrahydrofolate, Ahcy: adenosylhomocysteinase, BHMT: betaine homocysteine methyltransferase, MTases: methyltransferase, CBS: cystathionine-β-synthase, CTH: cystathionine-γ-lyase, MS: methionine synthase, SHMT: serine hydroxymethyltransferase. (**D**) Changes in the metabolites of the Met cycle after starvation of whole-body larvae. The SEM was calculated from six independent samples. Statistical significance was calculated using a two-tailed Student’s t-test: **p < 0.01, ***p < 0.001, and ****p < 0.0001. (**E**) Western blot analysis of Gnmt expression in the fat bodies of larvae after 24 h of starvation. αTub was used as a loading control. (**F**) Quantitative data of band intensity in (E). The SEM was calculated from four independent samples. Statistical significance was calculated using a two-tailed Student’s t-test; ** p < 0.01.

### Gnmt reduction due to inhibition of SAM production is controlled at the protein level, not the transcriptional level

In our previous study, the inhibition of SAM production in the FB of *Drosophila* larvae led to a decrease in Gnmt protein levels, which is highly expressed in the FB and utilizes SAM (7), suggesting the existence of a regulatory mechanism to decrease Gnmt levels and maintain SAM levels. Additionally, a decrease in Gnmt protein levels in the FB was observed during starvation in larvae (Fig. 1E and F). To visualize endogenous *gnmt* gene expression, we employed T2A-Gal4 (8), an expression driver that allows the manipulation of endogenous gene expression by combining a Gal4 driver with a self-cleaving T2A peptide, which induces ribosomal skipping during translation, enabling the simultaneous translation of two proteins. By inserting T2A-Gal4 into the 3’ end just before the stop codon of the *gnmt* gene using the clustered regularly interspaced short palindromic repeats (CRISPR)/Cas9 system, we generated *Drosophila* strains in which the Gal4 driver was inducible at the location of the endogenous *gnmt* gene (Fig. S1A). The expression of the fluorescent protein, GFP, specifically in *gnmt*-expressing cells allowed us to confirm its presence in the larval FB and detect the disappearance of the GFP signal upon *gnmt* knockdown (Fig. S1B). Notably, the GFP signal was not attenuated by *sams* knockdown, suggesting that the decrease in Gnmt due to inhibition of SAM production did not occur at the transcriptional level (Fig. S1B).

We confirmed the protein and gene expression levels of Gnmt by western blotting and quantitative PCR (qPCR), respectively, upon the knockdown of *sams* and *gnmt*. Knockdown of *sams* decreases the band of Gnmt, similar to *gnmt* knockdown. However, the amount of GFP, reflecting *gnmt* transcription induced by *Gnmt-T2A-Gal4*, did not decrease (Fig. 2A). In fact, the gene expression level of *gnmt* was not decreased by *sams* knockdown (Fig. 2B). These results suggest that the decrease in the SAM-consuming enzyme, Gnmt, due to inhibition of SAM production is controlled at the protein level, not the mRNA level. Additionally, we assessed the impact of *sams* and *gnmt* knockdown on SAM metabolism by performing metabolic measurements in the FB (Fig. S2). Knockdown of *sams* resulted in an increase in the levels of Met and a decrease in SAH, but SAM levels remained unchanged. Knockdown of *gnmt* resulted in a significant increase in SAM and MTA levels, indicating the crucial role of Gnmt in regulating their levels. These results support the hypothesis that SAM levels are maintained by decreased Gnmt levels during the inhibition of SAM production.

**Figure 2.**
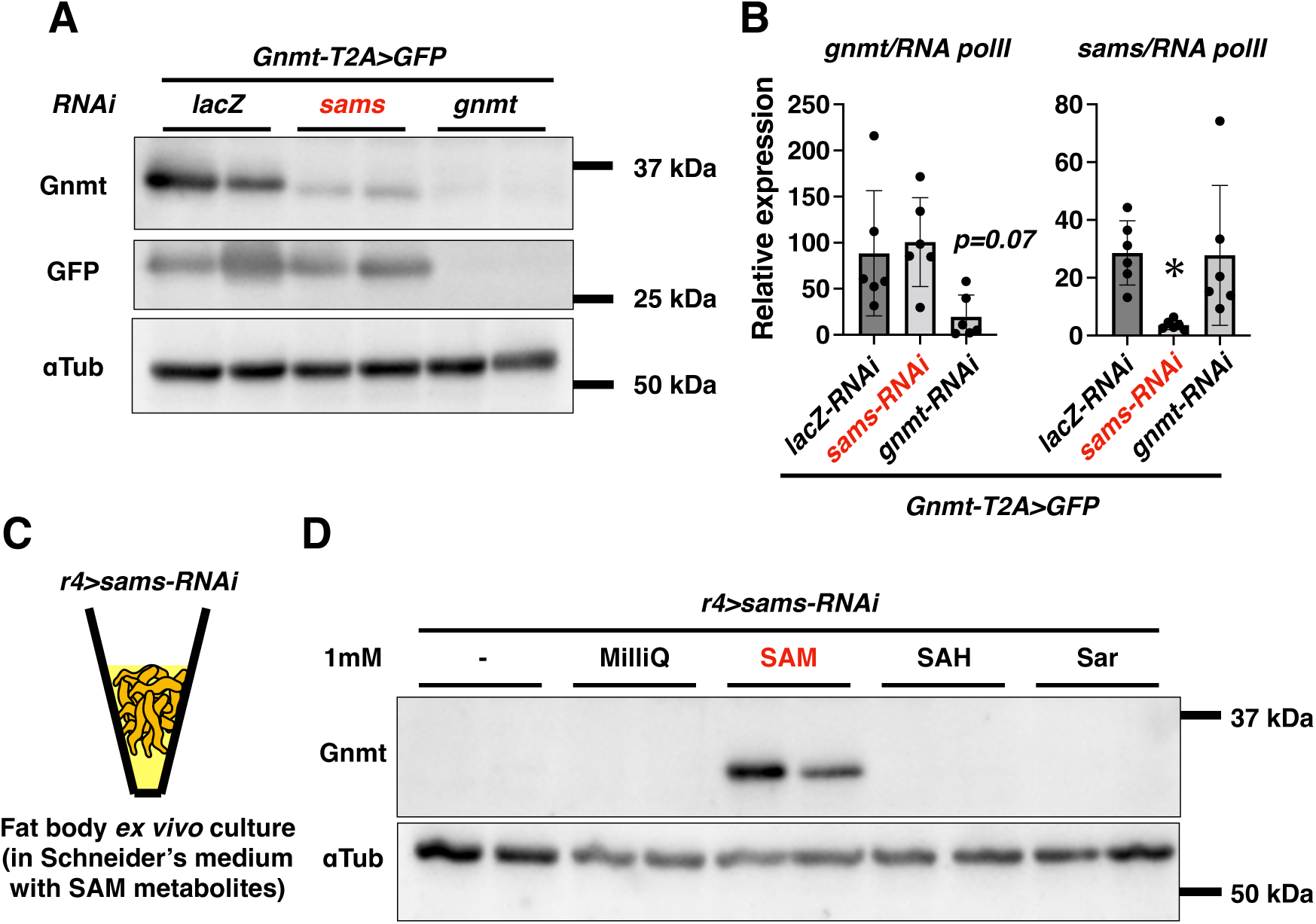
SAM buffering and Gnmt reduction in response to inhibition of SAM production. (**A**) Western blot analysis of Gnmt and GFP in the fat body of *Gnmt-T2A>GFP* larvae with knockdown of *lacZ*, *sams*, or *gnmt*. αTub was used as a loading control. Duplicate samples are indicated. (**B**) qRT-PCR analysis of *gnmt* and *sams* in the fat body of *Gnmt-T2A>GFP* larvae with genetic manipulation of *gnmt* or *sams* in the fat body. *RNA pol II* was used as an internal control. SEM was calculated from six independent samples. One-way ANOVA and Turkey’s multiple comparison test were employed to calculate significance: *p < 0.05. (**C**) Schematic of the fat body *ex vivo* culture in Schneider’s medium with SAM metabolites for 18 h. (**D**) Western blot analysis of Gnmt in the fat body of larvae with knockdown of *sams* and in each cultured condition. αTub was used as a loading control. Duplicate samples are indicated.

Furthermore, Met deprivation was induced using Methioninase, a Met-degrading enzyme (9). Induction of Methioninase in larval FB resulted in a global reduction in SAM metabolites and a decrease in Gnmt protein levels, not transcriptional levels (Fig. S3A-D).

### SAM administration rescues Gnmt decrease observed in FB *ex vivo* culture

To examine how the decrease in Gnmt induced by *sams-RNAi* was regulated by changes in SAM metabolism, we administered SAM metabolites associated with Gnmt to an *ex vivo* FB culture (Fig. 2C). The Gnmt decrease was rescued when 1 mM SAM was added to the medium (Fig. 2D). This rescue was also observed when 100 µM of SAM was added (Fig. S4A). Schneider’s Insect Medium contains 5.4 mM Met, suggesting that Gnmt was not rescued even at physiological Met levels. Additionally, measurement of SAM metabolic products in cultured FB revealed that administration of 100 µM SAM restored both SAM and MTA levels to those observed in control FB without *sams* knockdown (Fig. S4B). Furthermore, to test the possibility that the effect on Gnmt is due to less efficient uptake of SAH compared to SAM, we measured metabolite levels in *ex vivo* FB cultures supplemented with 100 µM SAH or Sar (Fig. S4C). No significant changes in Met or SAM levels were observed upon SAH or Sar supplementation. Notably, SAH levels increased within FB, indicating that SAH is taken up by FB. These results suggest that the observed effects are not simply due to differences in metabolite uptake by FB. Rather, it is suggested that SAM-dependent threshold exist for regulating Gnmt protein levels, and that the regulation of Gnmt is controlled by the sensing of this threshold.

### Ubiquitin-proteasome system (UPS) regulates the decrease in Gnmt levels and its nuclear accumulation

To elucidate the regulation of Gnmt protein levels, we focused on protein degradation pathways. To investigate the involvement of the major protein degradation pathways, autophagy, and the proteasome, we knocked down the components *Atg3* and *Rpn11* in FB. The decrease in Gnmt caused by *sams* knockdown was rescued by *Rpn11* knockdown but not by *Atg3* knockdown, revealing that the proteasome regulates the protein levels of Gnmt (Fig. 3A). Consequently, we attempted to identify the ubiquitin ligases that contribute to control. To identify key ubiquitin ligases, we analyzed our proteomic data from the FB (10)(The accession numbers of proteomics data are: PXD054313 for ProteomeXchange and JPST003071 for jPOST) and searched for ubiquitin-related factors specifically expressed in the FB. From Kyoto Encyclopedia of Genes and Genomes (KEGG) and Gene Ontology (GO) analyses, a total of 1112 proteins were identified. Of these nine KEGG terms for “Ubiquitin mediated proteolysis” and seven GO terms for “Ubiquitin ligase” were identified, resulting in a total of 11 factors after excluding duplicates. Small-scale RNAi screening was conducted for these 11 ubiquitin-related factors expressed in the FB. E1, E2, and E3 ubiquitin ligases were found to rescue Gnmt, similar to *Rpn11* knockdown (Fig. 3B). Although hyd and PPIL2 were identified as candidate E3 ligases, further verification with other RNAi strains confirmed that hyd consistently rescued Gnmt expression across multiple RNAi strains, confirming hyd as an E3 ligase (Fig. S5). HUWE1, a member of the HECT domain family, degrades PREX2, a phosphatase and tensin homolog (PTEN) inhibitor scaffolded by Gnmt (11, 12). Based on previous reports identifying HUWE1 as an E4 ligase (13), we conducted validation experiments and observed that knockdown of *HUWE1* rescued Gnmt (Fig. 3C). Thus, the decrease in Gnmt caused by *sams* knockdown was controlled by the proteasome and specific E1, E2, E3, and E4 ubiquitin ligases.

**Figure 3.**
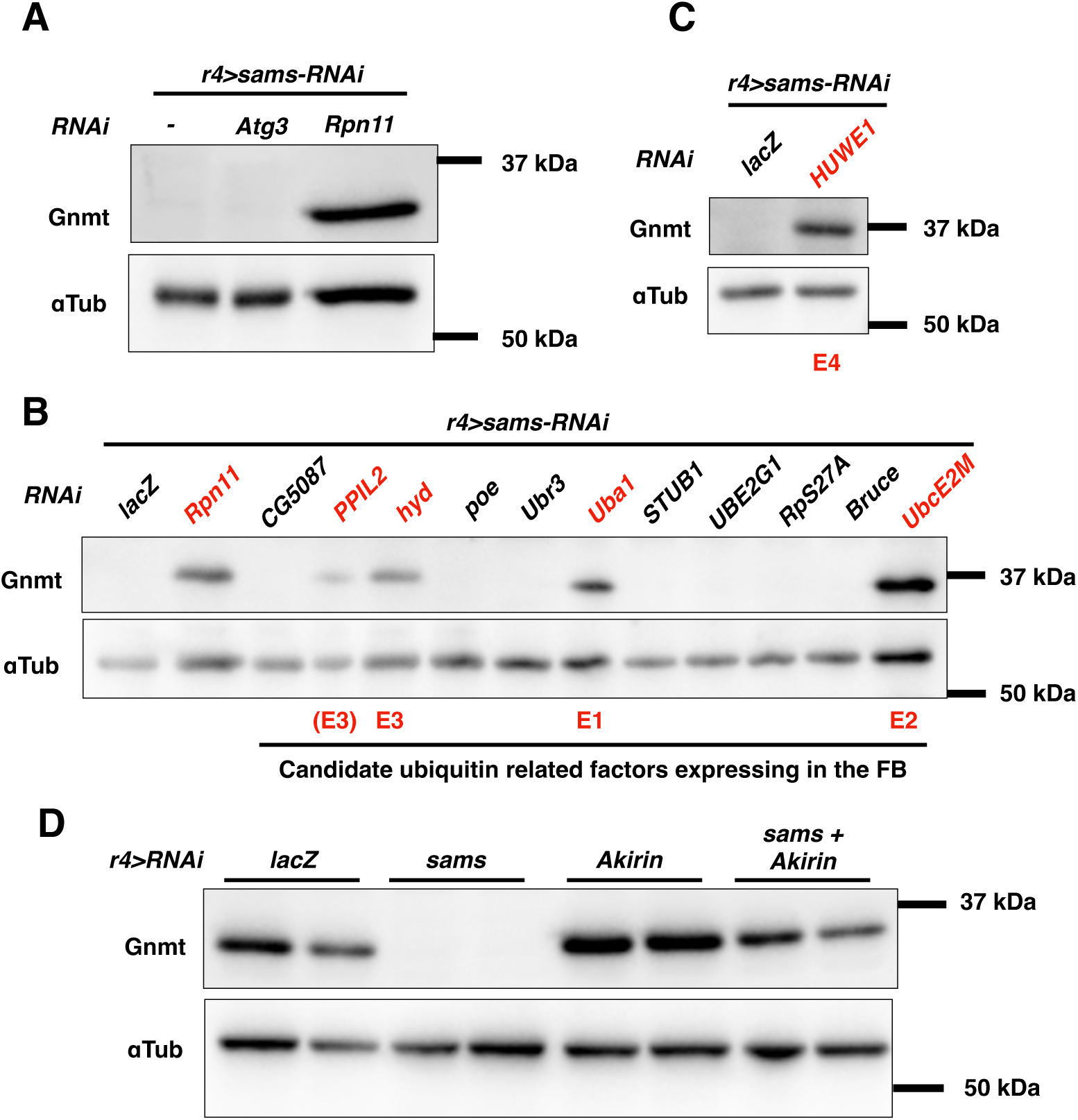
Regulation of Gnmt during inhibition of SAM production by the nuclear ubiquitin-proteasome system. (**A**) Western blot analysis of Gnmt in the fat body of larvae with knockdown of *Atg3* or *Rpn11*, (**B**) candidate ubiquitin-related factors expressed in the fat body, (**C**) *HUWE1*. αTub was used as a loading control. (**D**) Western blot analysis of Gnmt in the fat bodies of larvae with knockdown of *lacZ*, *sams,* or *Akirin*. αTub was used as a loading control. Duplicate samples are indicated.

### The decrease in Gnmt requires nuclear UPS, and its inhibition leads to Gnmt accumulation in the nucleus

While Gnmt is primarily localized in the cytosol, E3 and E4 ligases hyd and HUWE1, are localized in the nuclei of mammalian cells and *Drosophila* salivary glands, respectively (14, 15). Given the implications of the nuclear UPS in Gnmt regulation, attention has been shifted to the role of nuclear translocation of the proteasome. Using CRISPR screening in cultured mammalian cells has revealed that AKIRIN2 controls proteasome nuclear translocation (16). Consequently, knocking down *Akirin*, a *Drosophila* ortholog, an increase in Gnmt protein levels and rescue of the Gnmt decrease caused by *sams-RNAi* were observed (Fig. 3D).

Next, we investigated changes in Gnmt localization. We generated a *gnmt-V5-turboID* strain by inserting the V5 tag and the proximal labeling factor turboID at the C-terminus of *gnmt* (17). In this *gnmt-V5-turboID* strain, no significant differences in SAM metabolism status were observed compared to the wild-type (Fig. S6A). Additionally, when *gnmt* was knocked down using *r4-Gal4* driver, a notable increase in SAM was observed, similar to when *gnmt* was knocked down using *Gnmt-T2A-Gal4* driver without *gnmt-V5-turboID* (Fig. S6B and S2), suggesting functional integrity of the *gnmt-V5-turboID* knock-in strain. Furthermore, a decrease in Gnmt-V5-turboID detected by the V5 antibody upon *sams-RNAi* treatment was observed, similar to the decrease in Gnmt when *sams* was knocked down using *Gnmt-T2A-Gal4* (Fig. S6C, S6D, and 2A). Using this strain, we visualized changes in Gnmt induced by *sams*, *Akirin*, and *HUWE1* knockdown in the FB (Fig. 4A), and quantified the intensity of the V5 signal in both the whole cell and the nucleus of FB cells (Fig. 4B). Under normal conditions, the Gnmt-V5 signal was observed in both the cytosol and nucleus. However, while *Akirin-RNAi* led to an overall increase in intensity, the decrease in Gnmt caused by *sams-RNAi* resulted in its accumulation in the nucleus, specifically due to *Akirin-RNAi* (Fig. 4A and B). Similar changes were observed following *HUWE1* knockdown (Fig. 4A and B). To confirm these findings, additional RNAi lines for *HUWE1* and *Akirin* were examined. In all tested RNAi lines, the decrease in Gnmt induced by *sams-RNAi* was successfully reduced (Fig. S7A). Furthermore, the RNAi efficiency was validated by qPCR (Fig. S7B). These findings suggest that the nuclear proteasome and ubiquitin ligases regulate the stabilization of SAM levels by controlling the decrease in Gnmt levels upon inhibition of SAM production (Fig. 4F).

**Figure 4.**
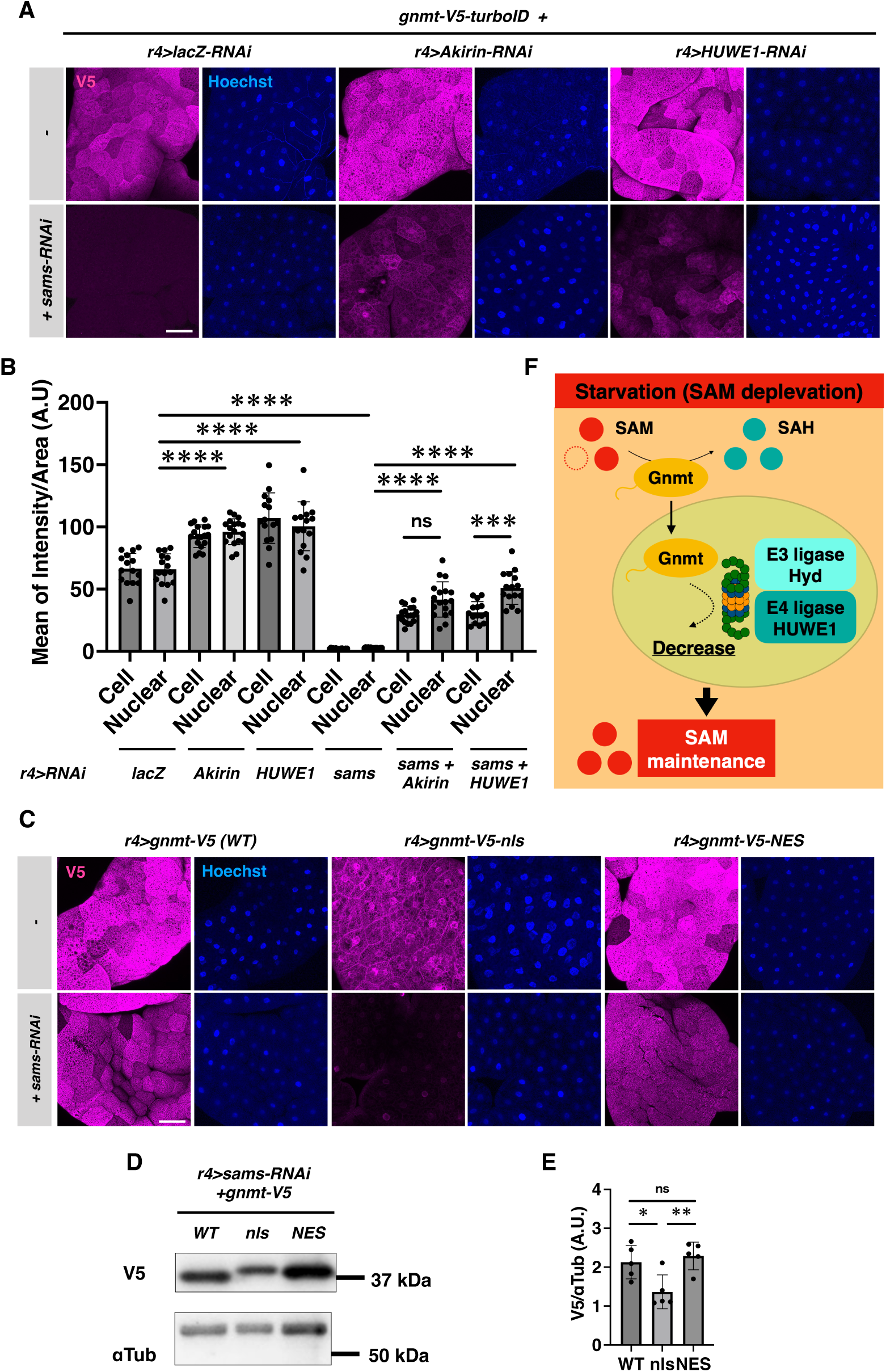
Regulation of Gnmt levels and nuclear localization by nuclear proteasome and ubiquitin ligase inhibition. (**A**) Representative images of larval fat body expressing *gnmt-V5-turboID* with knockdown of *lacZ*, *Akirin*, or *HUWE1*. The fat bodies were stained with anti-V5 and Hoechst. (Scale bar, 75 μm.) (**B**) Quantified V5 signal intensity in both the cell and nuclear area, as marked by Hoechest staining. n=15. Statistical significance was assessed using one-way ANOVA Turkey’s multiple comparison test was applied: ns, not significant; ***p < 0.001; ****p < 0.0001. (**C**) Representative images of larval fat body expressing *r4>gnmt-V5, -nls,* or *-NES* with or without knockdown of *sams*. The fat bodies were stained with anti-V5 and Hoechst. (Scale bar, 75 μm.) (**D**) Western blot analysis of Gnmt expression in the fat bodies of larvae expressing *r4>gnmt-V5, -nls,* or *-NES* with knockdown of *sams*. αTub was used as a loading control. (**E**) Quantitative data of band intensity in (D). The SEM was calculated from four independent samples. Statistical significance was assessed using one-way ANOVA Turkey’s multiple comparison test was applied: ns, not significant; *p < 0.05; **p < 0.01. (**F**) Schematic of SAM buffering and Gnmt decrease by nuclear UPS under starvation or inhibition of SAM production.

To demonstrate that nuclear translocation plays a key role in the regulation of Gnmt protein levels under *sams-RNAi* condition, we generated overexpression lines for Gnmt with added nuclear localization signal (nls) or nuclear export signal (NES). We constructed *UASz-gnmt-V5*, *UASz-gnmt-V5-nls*, and *UASz-gnmt-V5-NES* and performed tissue staining and western blotting (Fig. 4C-E). The reduced Gnmt levels in the Gnmt-V5-nls samples was detected by both tissue staining and western blotting. In contrast, no significant differences were observed between Gnmt-V5-NES and Gnmt-V5 (WT), which we attribute to Gnmt’s natural predominance in the cytoplasm. These findings support that Gnmt’s nuclear localization plays a significant role in regulating Gnmt protein levels.

Protein level of Gnmt is regulated by the detection of the SAM metabolism status (Fig. 2D), but the upstream mechanism remains unclear. SAMTOR and Unmet have been reported to function as SAM sensors upstream of the mTOR signaling pathway (18, 19). Therefore, we investigated whether knocking down these factors could rescue the decrease in Gnmt levels induced by *sams-RNAi*; however, this approach did not result in rescue (Fig. S8). This suggests that Gnmt regulation involves a mechanism that differs from those previously described.

We also attempted to detect Gnmt ubiquitination in larval FB. Using larval FB samples in which Gnmt degradation was impaired under *r4>sams-RNAi+Rpn11-RNAi* conditions, we performed immunoprecipitation (IP) of Gnmt, followed by IP-Western blotting using mono- and polyubiquitin antibodies (Fig. S9A). To enhance detection, we also added the deubiquitination inhibitor N-Ethylmaleimide (NEM). However, we could not detect ubiquitinated Gnmt.

Additionally, we employed an alternative approach to facilitate ubiquitin detection by adding GST-TR-TUBE (20) to FB lysates, which protects linear ubiquitin chains from deubiquitination. Despite these efforts, we still could not detect ubiquitinated Gnmt (Fig. S9B). The inability to detect Gnmt ubiquitination may be due to the inherently low abundance and instability of ubiquitinated proteins in vivo, as previously reported (21).

While it remains possible that Gnmt is not a direct ubiquitination target, our data suggest that nuclear E3 and E4 ubiquitin ligases, as well as Akirin, which facilitates nuclear translocation of the proteasome, are involved in Gnmt degradation (Fig. 3B-D). Furthermore, we also demonstrated that nuclear translocation plays a key role in the quantitative regulation of Gnmt (Fig. 4C-E). These findings indicate that nuclear UPS activity regulates Gnmt protein levels.

### Inhibition of HUWE1 in larval FB reduces lipid loss during starvation

Previous studies reported that *gnmt* mutant flies exhibit increased SAM levels and decreased starvation resistance (22). In addition, nicotinamide N-methyltransferase (NNMT), which produces 1-methylnicotinamide by catalyzing the methylation reaction with SAM and nicotinamide, has been proposed to function as a methylation sink in mouse white adipose tissue and liver, leading to SAM accumulation and promoting energy expenditure (23). To verify the effect of inhibition of Gnmt reduction by *HUWE1* knockdown under starvation conditions, we examined lipid storage and tolerance during starvation.

Starvation during the larval stage resulted in a decrease in Gnmt protein levels, as shown in Fig. 1E, but *HUWE1-RNAi* increased Gnmt levels and suppressed the starvation-induced decrease in Gnmt (Fig. 5A and B). Additionally, the amount of triglycerides (TAG) was increased by *HUWE1-RNAi,* which correlated with the amount of Gnmt (Fig. 5C). These results suggest that the suppression of Gnmt reduction by *HUWE1* knockdown affects lipid storage.

**Figure 5.**
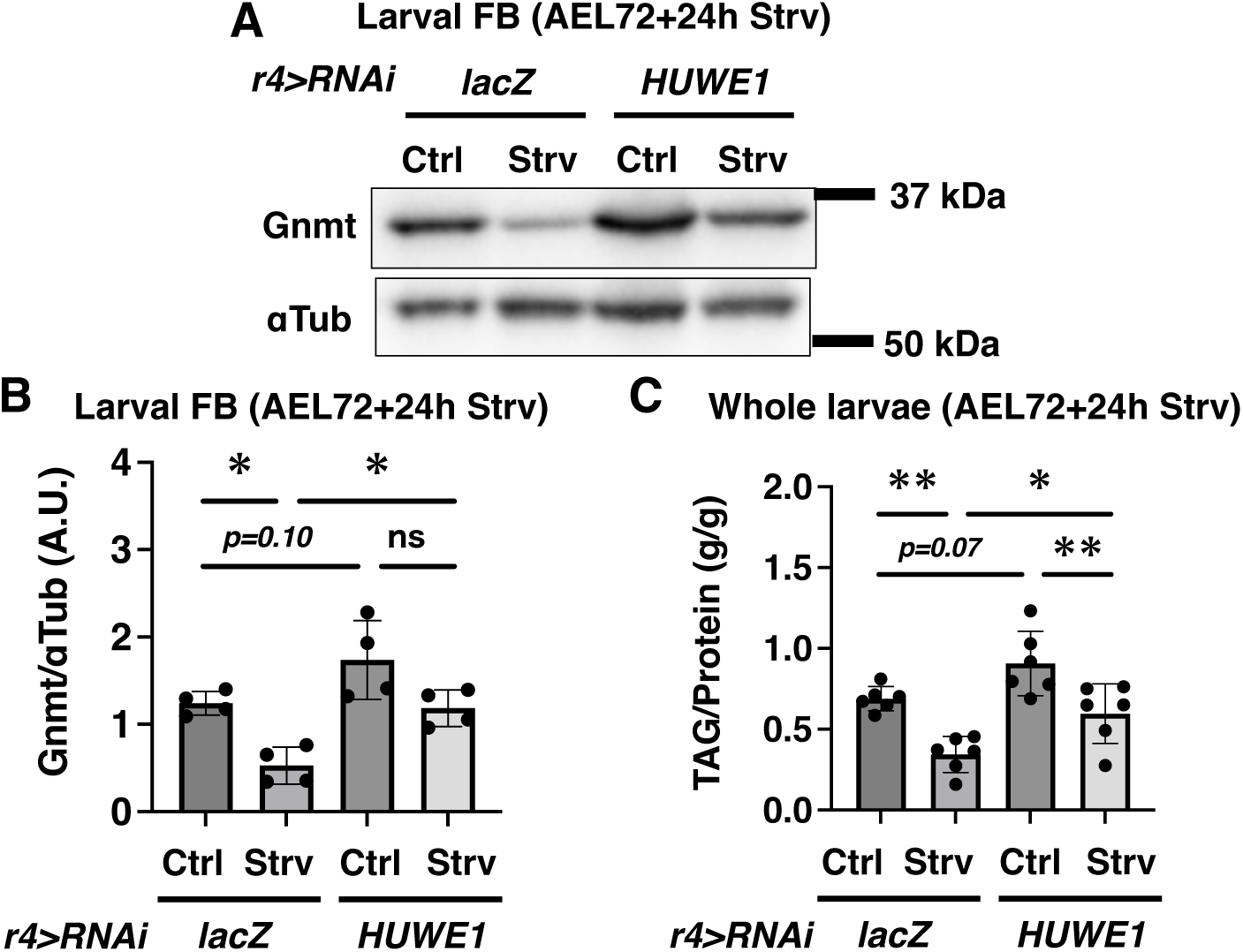
HUWE1 affects alteration of Gnmt and lipid storage caused by starvation in larval FB. (**A**) Western blot analysis of Gnmt in larval FB with FB-specific knockdown of *HUWE1* after 24 h starvation. αTub was used as a loading control. (**B**) Quantitative data of band intensity observed in (A). SEM was calculated from four independent samples. One-way ANOVA Turkey’s multiple comparison test was applied: ns, not significant; *p < 0.05. (**C**) Quantitative data on TAG levels in larvae. SEM was calculated from six independent samples. One-way ANOVA Turkey’s multiple comparison test was used for determining statistical significance: *p < 0.05; **p < 0.01.

We performed double knockdown of *HUWE1-RNAi* and *gnmt-RNAi* to assess the extent to which Gnmt contributes to TAG regulation as a HUWE1 target (Fig. S10). The trend of TAG reduction observed with *gnmt-RNAi* alone persisted in the double knockdown of *HUWE1-RNAi* and *gnmt-RNAi* (Fig. S10). This suggests that while HUWE1 has multiple targets, Gnmt is a major factor in TAG regulation.

## Discussion

In this study, the mechanism of SAM level stabilization mediated by the degradation of the SAM-utilizing methyltransferase Gnmt via the nuclear UPS was elucidated (Fig. 4F). The reduction in Gnmt by the nuclear UPS occurs under starvation and inhibition of SAM production (Fig. 1E and 2A). The inhibition of the Gnmt decreasing UPS machinery by knocking down *HUWE1* or *Akirin* in larvae leads to an increase in Gnmt levels. Consequently, suppression of lipid reduction was observed (Fig. 5). Gnmt is an enzyme that consumes SAM in the liver, adipose tissue and FB. The significance of Gnmt has been elucidated, including its role in energy expenditure associated with SAM accumulation, lifespan regulation, and regeneration of the *Drosophila* wing imaginal disc and liver (7, 22, 24–26). Our study revealed Gnmt as a major buffering system for SAM.

Rescue of Gnmt reduction by SAM administration in *ex vivo* FB cultures (Fig. 2D) suggested the presence of a SAM sensing system. However, known SAM sensors such as SAMTOR and Unmet did not rescue the decrease in Gnmt induced by *sams-RNAi* (Fig. S8). Recent reports indicate that Ubr1, a HECT domain family ligase similar to hyd and HUWE1, binds to amino acids during feeding, which triggers the degradation of its target Plin2, and promotes lipid storage stabilization (27). This implies that hyd and HUWE1 may contain a metabolite-sensing domain for the Met cycle or binding domains for SAM sensor proteins, serving as central hubs for metabolic sensing and degradation.

The reduction in Gnmt induced by *sams-RNAi* was rescued by inhibition of the nuclear UPS through the knockdown of *Akirin* and *HUWE1*, resulting in a notable increase in Gnmt accumulation in the nucleus (Fig. 4). Although Gnmt is predominantly localized in cytosol, carcinogen-induced nuclear translocation of GNMT has been reported in cultured mammalian cells (28). Protein kinase C-mediated phosphorylation of the N-terminus of GNMT was reported to contribute to this translocation, but the mechanism by which the SAM metabolic state influences GNMT nuclear translocation and the significance of this translocation remain unclear.

The *gnmt-V5-turboID Drosophila* line generated in this study enabled the exploration of the Gnmt proximal factors through biotinylation. By elucidating the metabolism-dependent changes in these factors, future research is expected to identify the mechanisms underlying nuclear translocation and the sensing mechanism of SAM.

The degradation of metabolic enzymes was shown to be regulated by the E3 ligase UBR5, an ortholog of hyd. Under high glucose conditions, acetylated phosphoenolpyruvate carboxykinase interacts with UBR5 and undergoes degradation (29). In our study, the control of Gnmt degradation via hyd was elucidated, suggesting a SAM level-dependent interaction between Gnmt and hyd.

In this study, the degradation of Gnmt by the nuclear UPS was important for SAM buffering, while its inhibition led to an increase in Gnmt and TAG in larvae (Fig. 5). We have previously reported that Gnmt expression is induced by Toll/dFOXO when tissue necrosis is induced by apoptosis inhibition in adult flies (22). *Gnmt* mutant led to a reduction in neutral fat TAG and was sensitive to starvation, suggesting that the decrease in SAM levels due to Gnmt elevation is a protective response against energy expenditure under inflammatory conditions.

Additionally, in AML12 cells, a mouse liver cell line expressing the SAM-consuming enzyme NNMT, the inhibition of NNMT resulted in SAM accumulation and a decrease in neutral fat (30). Moreover, knockdown of *Nnmt* in mouse white adipose tissue and liver enhanced energy expenditure and protected against diet-induced obesity (23). Additionally, in *Nnmt* knockout mice, SAM accumulates in the liver upon the transplantation of cancer cells, alleviating the inhibition of the urea cycle by cancer cells (31). These results suggest that the regulatory mechanisms of SAM-consuming MTases, such as NNMT and Gnmt, are conserved across species and play crucial roles in energy expenditure and the response to starvation. Understanding the regulatory mechanisms for SAM buffering will provide us with unidentified insights into the regulation of SAM-related physiologies and pathologies.

## Materials and Methods

Complete materials and methods used in this study are described in *SI Appendix, Materials and Methods.* Sample preparation and antibody staining are detailed in *SI Appendix, Materials and Methods*. *Gnmt-T2A-Gal4* and *gnmt-V5-turboID* knock-in flies were generated using CRISPR/Cas9. Metabolites were analyzed using a SHIMADZU LCMS-8060 (Kyoto, Japan). The sample preparation and procedures for the starvation assay, western blotting, immunohistochemistry, quantitative RT-PCR, and metabolite analyses are detailed in *SI Appendix, Materials and Methods*.

## Author Contributions

S. K. and M. M. conceived the study. S. K. and M. M. designed the experiments and wrote the manuscript. S. K. performed the experiments and analyzed the data. M. M. supervised the study.

## List of Abbreviations

FB: fat body
Gnmt: glycine N-methyltransferase
GO: Gene Ontology
Hcy: homocysteine
KEGG: Kyoto Encyclopedia of Genes and Genomes
Met: methionine
MTA: methylthioadenosine
NNMT: nicotinamide N-methyltransferase
qPCR: quantitative PCR
SAH: S-adenosylhomocysteine
SAM: S-adenosylmethionine
Sar: Sarcosine
Sams: SAM synthetase
TAG: triglycerides
TSF: transsulfuration
UPS: ubiquitin-proteasome system
UTR: untranslated region

## Acknowledgments

We thank the Bloomington *Drosophila* Stock Center and the Vienna *Drosophila* Resource Center for the fly stocks. We are grateful to S. Kondo (Tokyo University of Science) for the pBluescript II SK(+) and pPGxRF3 vectors, to N. Perrimon (Harvard Medical School) for *UAS-Methioninase*, and to F. Ikeda (Osaka University) for the pGEX6p1-GST-TR-TUBE vector. We also thank S. Murata for valuable insights into the nuclear UPS. This work was supported by grants from AMED-Project for Elucidating and Controlling Mechanisms of Aging and Longevity (Grant Number. JP21gm5010001 to M.M.). Additional support was provided by the Japan Society for the Promotion of Science to M.M. (Grant Numbers. 21H04774, 23H04766, and 24H00567) and to S.K. (Grant Numbers 21K15100 and 24K09774). This research was also supported by the Japan Science and Technology Agency (JST), PRESTO, under Grant Number JPMJPR24N4 to S.K., and by a grant from the Takeda Science Foundation to S.K.

We thank the members of the Miura laboratory for their technical assistance and suggestions, especially N. Shinoda for providing the EGFP antibody, K. Takenaga for preparing the fly food, A. F. Kashio for constructing the pUASz-gnmt-V5 vectors, and Y. Yoshida for the proteomic data. We also appreciate the helpful discussions and comments from N. Shinoda and Y. Nakajima.

