## Supplemental Information for "S-adenosylmethionine metabolism buffering is regulated by glycine N-methyltransferase decrease via nuclear ubiquitin-proteasome system"

[a] Department of Genetics, Graduate School of Pharmaceutical Sciences, University of Tokyo, 7-3-1 Hongo, Bunkyo-ku, Tokyo 113-0033, Japan; [b] Department of Integrative Bioanalytics, Institute of Development, Aging and Cancer (IDAC), Tohoku University, Sendai 980-8575, Japan; [c] Laboratory for Cell Vigor Regulation, National Institute for Basic Biology, Nishigonaka 38, Okazaki, Aichi 444-8585, Japan

[1] Corresponding authors: Soshiro Kashio, Masayuki Miura

**Supporting Information**

### Materials and Methods

#### Fly strains and genetics

*r4-Gal4*, *UAS-GFP*, *UAS-gnmt-RNAi*, and *UAS-lacZ-RNAi* have been previously characterized (7, 26). *UAS-Rpn11-RNAi* (#33662), *UAS-HUWE1-RNAi HMS1* (#36714), *UAS-HUWE1-RNAi HMS2* (#36715), *UAS-Akirin-RNAi HMS* (#34036), *UAS-hyd-RNAi* (#32352), *UAS-poe-RNAi* (#32945), *UAS-Uba1-RNAi* (#76066), *UAS-STUB1-RNAi* (#34017), *UAS-UBE2G1-RNAi* (#64032), and *UAS-PPIL2-RNAi* (#56975) were obtained from the Bloomington *Drosophila* Stock Center (Bloomington, IN, USA). *UAS-sams-RNAi 1* (v103143), *UAS-sams-RNAi 2* (v7168), *UAS-samDC-RNAi* (v101753), *UAS-dph1-RNAi* (v103550), *UAS-dph5-RNAi* (v110771), *UAS-Rpn11-RNAi* (v19272), *UAS-CG5087-RNAi* (v106663), *UAS-PPIL2-RNAi* (v105644), *UAS-hyd-RNAi* (v44676), *UAS-Ubr3-RNAi* (v106993), *UAS-RpS27A-RNAi* (v105501), *UAS-Bruce-RNAi* (v107620), *UAS-UbcE2M-RNAi* (v100761), *UAS-HUWE1-RNAi* (v330156), *UAS-Akirin-RNAi* (v109671), and *UAS-Atg3-RNAi* (v101364) were procured from the Vienna Drosophila Resource Center (Vienna, Austria). *CantonS* was used as the wild type. *UAS-Methioninase* (9) was provided by Dr. Nobert Perrimon. Images of the larvae were captured using a Leica microscope MZ10F.

#### Starvation assay

For starvation studies, larvae were collected at the early third instar stage (AEL 72 h) and transferred to 1% agar (Kishida Chemical)/MilliQ. The larval samples were collected after 24 h of incubation. Statistical analyses and data visualization were conducted using the RStudio software (ver. 2023.09.1+494).

#### Western Blotting

The fat bodies of larval were dissected in phosphate-buffered saline (PBS) and homogenized in radio-immunoprecipitation assay buffer with a protease inhibitor cocktail (Roche). Samples were boiled for 5 min at 98°C after mixing with 6 × Laemmli sample buffer (1 M Tris-HCl, pH 6.8), 12% sodium dodecyl sulfate (SDS), 0.6% bromophenol blue, 15% 2-mercaptoethanol). Approximately 10 µg protein was subjected to SDS-polyacrylamide gel electrophoresis and transferred onto a polyvinylidene difluoride membrane (Millipore). After 30 min blocking with 4% skim milk, membranes were incubated with the primary antibody at 4°C overnight. Representative images from at least two independent experiments with reproducible results are presented. Anti-EGFP polyclonal antibodies were generated by Shinoda using the QSALSKDPNEKRDH peptide as an antigen and were produced by Eurofins, Inc. The antibodies and dilutions used are listed in the following table.

| | Antibody or Probe | Host | Origin | Cat. number | Dilution |
|---|---|---|---|---|---|
| 1st antibody | anti-Gnmt | Rabbit | Ref. 22 | NA | 1/2000 |
| | anti-V5 | Mouse | Invitrogen | R96025 | 1/2000 |
| | anti-alpha Tubulin | Mouse | Cell Signaling Technology | 3873S | 1/2000 |
| | anti-EGFP | Rabbit | N. Shinoda | NA | 1/1000 |
| | anti-Ubiquitin | Mouse | Stressgen | SPA-203 | 1/1000 |
| 2nd antibody | Anti-Mouse IgG HRP Conjugate | Goat | Promega | W4021 | 1/2000 |
| | Anti-rabbit IgG, HRP-linked Antibody | Goat | Cell Signaling Technology | 7074S | 1/2000 |
| | Streptavidin-HRP Conjugate | | Invitrogen | SA10001 | 1/2000 |

#### Immunohistochemistry

Larval tissues were dissected in PBS, fixed in 4% paraformaldehyde, and washed with 0.3% Triton X-100 in PBS. The antibodies, probes, and dilutions used are listed in the following table. Confocal images were captured using Leica SP8.
For quantification of the V5 signal in Fig. 4B, the entire cell or the region of the nucleus indicated by Hoechst staining was quantified using Fiji software.

| | Antibody or Probe | Host | Origin | Cat. number | Dilution |
|---|---|---|---|---|---|
| 1st antibody | anti-V5 | Mouse | Invitrogen | R96025 | 1/500 |
| 2nd antibody | Anti-mouse-IgG, 647 | Donkey | Thermo Fisher Science | A-31571 | 1/500 |
| Probe | Hoechst33342 | | Invitrogen | H3570 | 0.8 µM |
| | Streptavidin-Cy2 | | Jackson Immuno Research | 016-220-084 | 1/500 |

**Quantitative RT-PCR**

The total RNA from five larval fat bodies and four whole larvae was extracted using the ReliaPrep RNA Tissue Miniprep System (Promega). Total RNA (200ng) was used for cDNA synthesis using the PrimeScript RT Reagent Kit with gDNA Eraser (TAKARA). Quantitative PCR was performed using TAKARA TB Green Premix Ex Taq II (Tli RNase H Plus) on a QuantStudio 6 Real-Time PCR system (Thermo Fisher). *RNA pol II* was used as the internal control. The primer sequences are listed in the following table.

| | Forward (5' ~3' ) | Reverse (5' ~3' ) |
|---|---|---|
| *RNA polII* | CCTTCAGGAGTACGGCTATCATCT | CCAGGAAGACCTGAGCATTAATCT |
| *gnmt* | CCGCTGACAGTGTGTTCGTT | CACGCTTGCATCCCTTATTGC |
| *sams* | AATGTGCGACCAAATCAGCG | TTCGCGTTTGGATCCTGCTT |
| *HUWE1* | CGAGCTTATTGATGAGCTGACC | GCAGTAGTGTGACATCCGTTTC |
| *Akirin* | TGTGCAACCCTGAAACGAG | ACGGACTAGGTTCGGTGCTAT |

**Generation of *Gnmt-T2A-Gal4* knock-in fly using CRISPR/Cas9 system**

*Gnmt-T2A-Gal4* knock-in flies were generated as described previously (8). Clustered regularly interspaced short palindromic repeats (CRISPR)/Cas9 genome editing was used to create a *Gnmt-T2A-Gal4* knock-in fly with a T2A-Gal4 sequence inserted immediately before the end codon of *gnmt*. Double-strand breaks were generated by the sgRNA-Cas9 protein at two locations. Consequently, following homologous recombination repair of the break site, the DNA sequence with the fluorescent marker protein 3xP3-RFP was inserted instead, containing two break sites: one approximately 200 bases (homology arm) and the other sequence containing 3xP3-RFP, which were incorporated into the donor vector pBluescript II SK(+). The donor and sgRNA expression vectors were introduced into transgenic lines expressing Cas9 protein under the control of the Act5C promoter. *Gnmt-T2A-Gal4* knock-in flies were selected using 3xP3-RFP expression as an indicator.
Pairs of guide RNAs were identified using the CRISPR Optimal Target Finder tool available on flyCRISPR website (http://flycrispr.molbio.wisc.edu/). The DNA fragments for the guide RNAs were subcloned using a DNA Ligation Kit (TAKARA) into the *Bbs*I-digested U6b-sgRNA-short vector (a gift from N. Perrimon). Primers listed in the following table were annealed to generate DNA fragments for guide RNAs.

| | Forward (5' ~3' ) | Reverse (5' ~3' ) |
|---|---|---|
| Target 1 | TTCGGATCAGGTGAATGTAGAAGG | AAACCCTTCTACATTCACCTGATC |
| Target 2 | TTCGGCCAAAGCACAGGGTCACCA | AAACTGGTGACCCTGTGCTTTGGC |

DNA fragments of the 5′ and the 3′ homology arms were amplified from fly genomic DNA. The T2A-Gal4-3xP3-RFP cassette DNA fragment was amplified from pPGxRF3. The three fragments were assembled by extension PCR using PrimeStar Max DNA polymerase (TAKARA) and cloned into linearized pBluescript II SK(+) with *Eco*RI using In-Fusion HD Cloning (TAKARA).
The PAM sequences of the gRNA-binding sites in the donor template were mutated to prevent Cas9-directed cleavage following the homology-directed repair. The DNA fragments were amplified by PCR using the following primers:

| | Forward (5'~3') | Reverse (5'~3'') |
|---|---|---|
| 5' Homology Arm | TCGCTGAAGGTCTCCGCTCTTCTACTGCGG | CTACTGCAGCTTTTCGGCAGACCGC |
| 3' Homology Arm | TCTTTCTAGGGTTAATCGCGGCCTCGATCAC | ATTGACGGCTCTTCAAATGGAAAAGCATTTGC |

Each RFP-positive transformant was isogenized and verified through genomic PCR and sequencing. The 3xP3-RFP marker was removed by crossing with the TM6B hs-Cre line (Bloomington #1501), which exhibited a leaky expression. Subsequently, RFP-negative F2 flies were collected.

**Generation of *gnmt-V5-turboID* knock-in fly using CRISPR/Cas9 system**
*gnmt-V5-turboID* knock-in flies were generated as described previously (17). CRISPR/Cas9 genome editing was used to create *gnmt-V5-turboID* knock-in fly with a V5-turboID sequence inserted immediately before the end codon of *gnmt*. Double-strand breaks were generated by the sgRNA-Cas9 protein at two locations. Consequently, following homologous recombination repair of the break site, a DNA sequence with the fluorescent marker protein DsRed was inserted containing two break sites: one sequence approximately 200 bases (homology arm) and the other sequence containing DsRed in between, which were incorporated into the donor vector pBac. Donor and sgRNA expression vectors were introduced into transgenic lines expressing the Cas9 protein under the control of the Act5C promoter. The *gnmt-V5-turboID* knock-in flies were selected using DsRed expression as an indicator.
Pairs of guide RNAs were identified using the CRISPR Optimal Target Finder tool available on flyCRISPR (http://flycrispr.molbio.wisc.edu/). The DNA fragments for the guide RNAs were subcloned using a DNA Ligation Kit (TAKARA) into the *Bbs*I-digested U6b-sgRNA-short vector (a gift from N. Perrimon). Primers listed in the following table were annealed to generate DNA fragments for guide RNAs.

| | Forward (5'~3') | Reverse (5'~3') |
|---|---|---|
| Target 1 | TTCGGATCAGGTGAATGTAGAAGG | AAACCCTTCTACATTCACCTGATC |
| Target 2 | TTCGGCCAAAGCACAGGGTCACCA | AAACTGGTGACCCTGTGCTTTGGC |

To generate the homology-directed repair template, DNA fragments of the 5′ and the 3′ homology arms were assembled into the *Bsa*I site of pBac[3xP3-DsRed_polyA_Scarless_TK] (generated by T. Katsuyama) using NEBuilder HiFi DNA Assembly (New England Biolabs) or In-Fusion HD Cloning (TAKARA).
The PAM sequences of the gRNA-binding sites in the donor template were mutated to prevent Cas9-directed cleavage following the homology-directed repair. The DNA fragments were PCR amplified using the following primers:

| | Forward (5' ~3' ) | Reverse (5' ~3' ) |
|---|---|---|
| 5' Homology Arm | ATAAGCTTGATATCGTCCGTGCTCTTCTACTGCGG | GCTGCCTCTGCCCTCAACCTGTGGCTTTTCGATCAGGTGAATGTAGAAGGCCGG |

| | | |
|---|---|---|
| 3' Homology Arm | CGGCCTCGATCACGAACGGG | CGGGCTGCAGGAATTACGTAATGGAAAAGCATTTG |

Each DsRed-positive transformant was isogenized and verified through genomic PCR and sequencing. The detailed plasmid DNA sequences are available upon request. Transgenic flies were generated by BestGene, Inc.

**Metabolites extraction from larvael samples**

Metabolites from whole bodies or tissue samples were extracted in 80% methanol from frozen 5 whole flies or larvae and five larval fat bodies. Samples were deproteinized with 50% acetonitrile. The supernatant was collected, transferred to a prewashed centrifugal filter unit (Nanosep with 10 K Omega, Pall Corporation), and centrifuged at 14,000 × *g*, 4°C for 10–30 min. The contents of the sample tubes were evaporated using a centrifugal concentrator CC-105 (TOMY). The pellets were dissolved in Milli-Q water and stored at -80°C until analysis.

The amount of protein was quantified for normalization. Briefly, proteins were extracted from the precipitate remaining after the metabolite extraction. The precipitates were washed in acetone, and the protein was extracted using 0.1 N NaOH at 95 °C for 5 min. Protein content was measured using a BCA protein assay kit (Thermo Fisher Scientific).

**UPLC-MS/MS**

The metabolites were measured using ultra-high-performance liquid chromatography equipped with tandem mass spectrometry (SHIMAZU 8060) based on the Primary Metabolites package ver.2 (Shimadzu) (26). For metabolite analysis, the extracted samples were injected to UPLC-MS/MS with a PFPP column (Discovery HS F5 (2.1 mm × 150 mm, 3 µm), Sigma–Aldrich) in a column oven at 40 °C. A gradient from solvent A (0.1% formic acid, water) to solvent B (0.1% formic acid, acetonitrile) was performed for 20 minutes. The concentration of the metabolites was determined using a standard curve obtained from serial dilutions of the standard solution for each metabolite, and parameter optimization with the use of software (Labsolutions, Shimadzu). Statistical analyses were conducted using the GraphPad Prism 9 or R Studio software (ver. 2023.09.1+494).

**Triglyceride (TAG) measurement assay.**

TAG levels in larvae and adult males were measured using a LabAssay™ Triglyceride Assay Kit (WAKO). Briefly, five larvae or five adult males were collected in a 1.5 mL tube and snap-frozen in liquid nitrogen. Each sample was extracted using cold PBST (5% Triton-X in PBS) and prepared for colorimetric assays using the kit. TAG absorbance was measured at 595 nm with ARVOx3 (PerkinElmer) and data were adjusted to the total TAG level per total protein content (measured using a BCA protein assay kit [Thermo Fisher Scientific]).

**Generation of *UAS-gnmt-V5*, *UAS-gnmt-V5-nls*, *UAS-gnmt-V5-NES* transgenic flies**

The sequences of *gnmt-V5, gnmt-V5-nls, gnmt-V5-NES* were PCR-amplified from *pUASz-gnmt-V5-turboID* vector. The sequences of nls and NES were as follows; nls: ccaaagaaaaagagaaaagta, NES: ctgcctcccctggagcgcctgaccctg. The amplified fragments were inserted into *pUASz1.0* (DGRC, #1431) vector with *Xho*I and *BamH*I restriction using In-Fusion HD Cloning (TAKARA). This sequence was confirmed by Eurofins, Inc. The vectors were injected into the transgenic *y1 w67c23; P{CaryP}attP40* strain. Transgenic flies were generated by Best Gene Inc. Each red eye-positive transformant was isogenized.

**Co-immunoprecipitation**

Larval fat bodies were lysed on ice using IP lysis buffer [25 mM Tris–HCl (pH 7.5), 100 mM NaCl, 2 mM EDTA, 0.5% Triton X-100] containing complete EDTA-free protease inhibitor cocktail. The samples were adjusted to 440 µg protein/440 µL IP lysis buffer. The lysates were centrifuged at 20 000 ***g***, 4 °C for 5 min. Supernatants (40 µL) were collected as input. The remainder of the supernatant was incubated overnight at 4 °C with 20 µL anti-V5-tag mAb-Magnetic Agarose (#M167-10, MBL, Minato-ku, Japan) equilibrated with the IP lysis buffer. The beads were washed thrice in washing buffer [50 mM Tris-HCl (pH 7.5), 500 mM NaCl, 0.1% NP40, and 0.05% sodium deoxycholate] and boiled for 5 min at 95 °C with 50 µL

1× Laemmli buffer. The samples were magnetically separated, and the supernatants were subjected to SDS/PAGE.

**GST-TR-TUBE pulldown assay**

GST-TR-TUBE proteins were expressed by *pGEX6p1-GST-TR-TUBE* vector in *E. coli* BL21 (20). 1mM IPTG was added and cultured overnight at 16°C. The bacteria were harvested by centrifugation and resuspended in GST buffer (20 mM Tris, pH 8.0, 100 mM NaCl) supplemented with complete protease inhibitors (Roche Diagnostics), and sonicated. The cleared lysate was incubated with GST-accept beads (Nacalai Tesque) for 2 h at 4 °C with continuous rolling. After the beads were washed three times with PBS, proteins were eluted with 10 mM glutathione in 50 mM Tris-HCl buffer, pH 8.0.

Larval fat bodies were lysed with chilled lysis buffer (50 mM Tris–HCl, pH 7.5, 100 mM NaCl, 5 mM MgCl2, 10% glycerol, 0.2% NP-40, 1 mM PMSF, 20 mM NEM, and Complete protease inhibitors). Cleared lysates were subjected to GST pulldown at 4°C overnight using GST-TR-TUBE. After washing with chilled lysis buffer for five times, the pulldown samples were subjected to SDS–PAGE followed by immunoblotting. Input of the GST-TR-TUBE was analyzed by CBB staining.

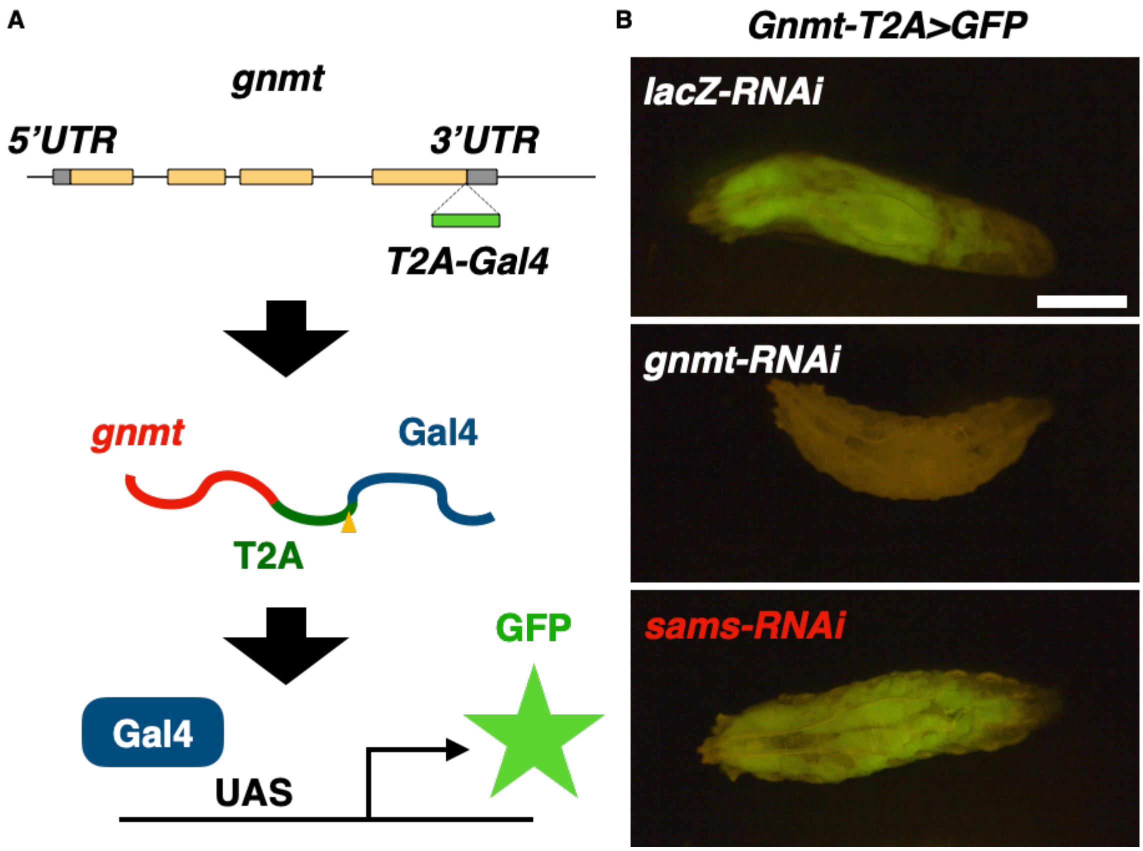


**Fig. S1. Visualization of *gnmt* expression by *Gnmt-T2A-Gal4*.**

(A) Schematics of *Gnmt-T2A-Gal4* driver and GFP induction. Gal4 and the T2A peptide were knocked in immediately before the stop codon. Gal4 was expressed with endogenous *gnmt* and the self-cleaved peptide T2A (the yellow arrowhead indicates the self-cleavage site). (B) Representative images of *Gnmt-T2A>GFP* expressing larvae after the genetic manipulation of *sams* or *gnmt*. (Scale bar, 1 mm.)

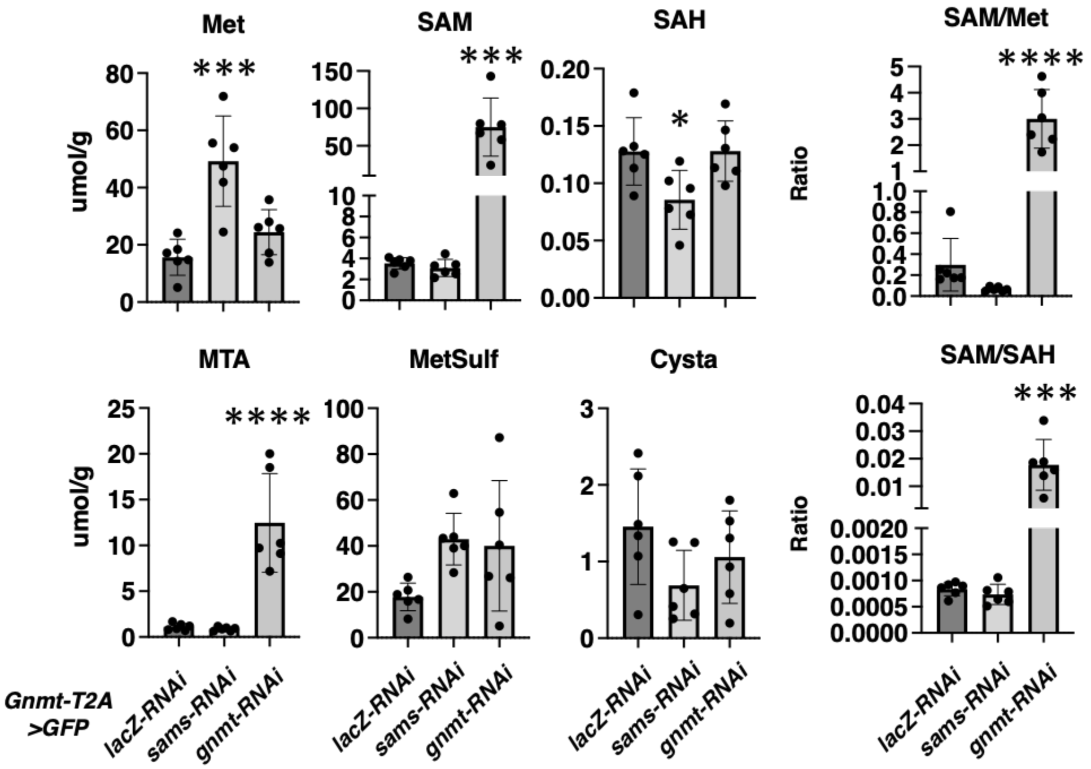


**Fig. S2. Metabolic changes of SAM metabolism by knockdown of enzymes.**

Changes in metabolite levels of SAM metabolism by genetic manipulation of *gnmt* or *sams* in the fat body. Metabolites were extracted from the whole larvae. The SEM was calculated from six independent samples. Statistical significance assessed using one-way ANOVA Tukey’s multiple comparison test was used: *p < 0.05, ***p < 0.001, and ****p < 0.0001.

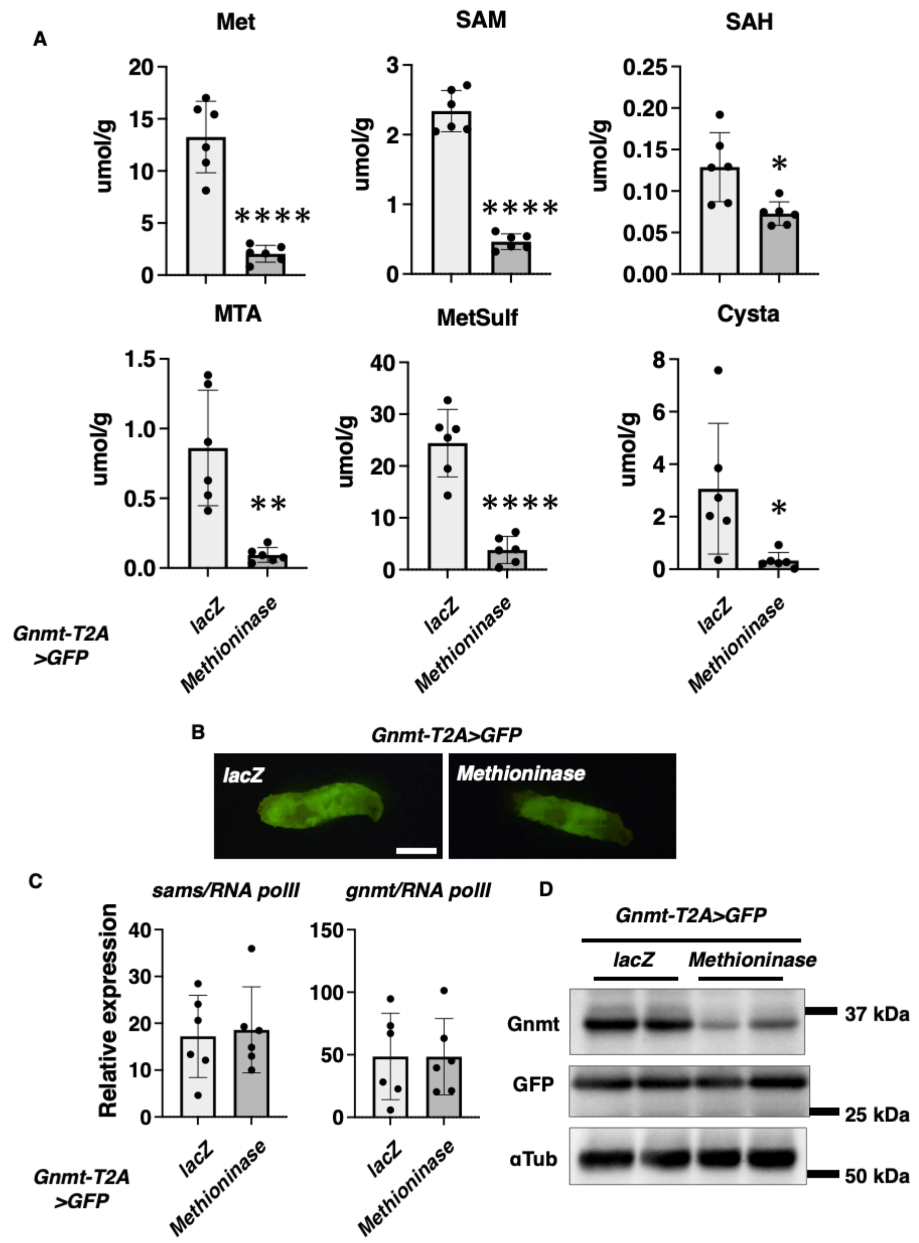


**Fig. S3. Met deprivation-induced decrease in Gnmt at the protein level.**

(A) The metabolites level of Met metabolism changes by genetic Met restriction with overexpression of *Methioninase* in the fat body. Metabolites were extracted from whole larvae. SEM was calculated from six independent samples. Statistical significance was calculated using the two-tailed student's t-tests: *$p < 0.05$; **$p < 0.01$; ****$p < 0.0001$. (B) Representative image of *Gnmt-T2A>GFP* expressing larvae with genetic overexpression of *Methioninase*. (Scale bar, 1 mm.) (C) qRT-PCR analysis of *gnmt* and *sams* in the fat body of *Gnmt-T2A>GFP* expressing larvae with genetic overexpression of *Methioninase* in the fat body. *RNA polII* was used as an internal control. (D) Western blot analysis of Gnmt and GFP in the fat body of *Gnmt-T2A>GFP* expressing larvae with genetic overexpression of *Methioninase*. αTub was used as a loading control. Duplicate samples are indicated.

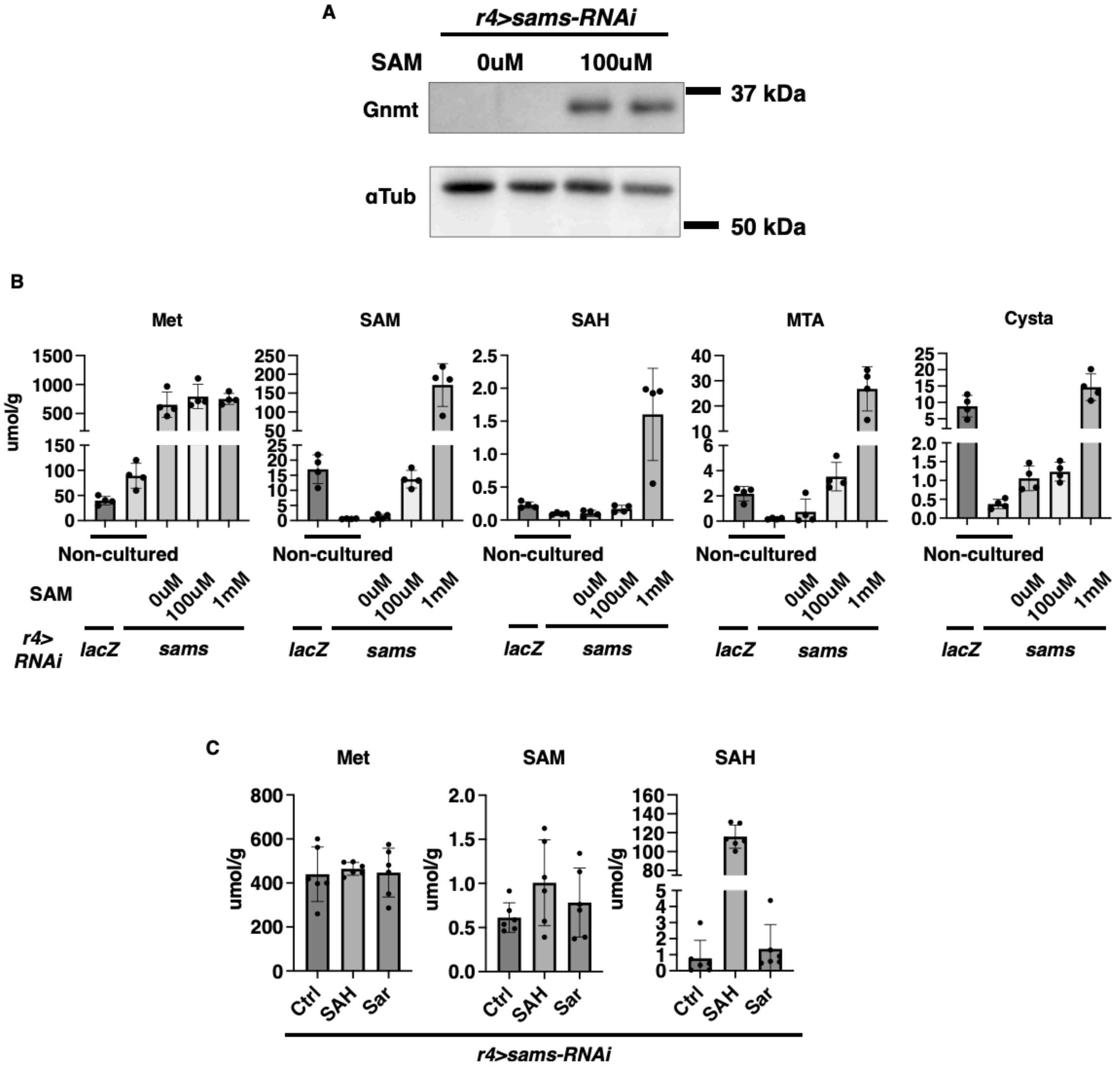


**Fig. S4. Gnmt rescue and SAM metabolic changes induced by SAM administration in FB *ex vivo* culture.**

(A) Western blot analysis of Gnmt in the fat body of larvae with knockdown of *sams* and in different cultured conditions. αTub was used as a loading control. Duplicate samples are indicated. (B) The changes in metabolite levels of SAM metabolism in the fat body after different cultured or non-cultured condition. SEM was calculated from four independent samples. (C) The metabolites level of SAM metabolism changes in the fat body after each cultured condition (100uM). SEM was calculated from four independent samples.

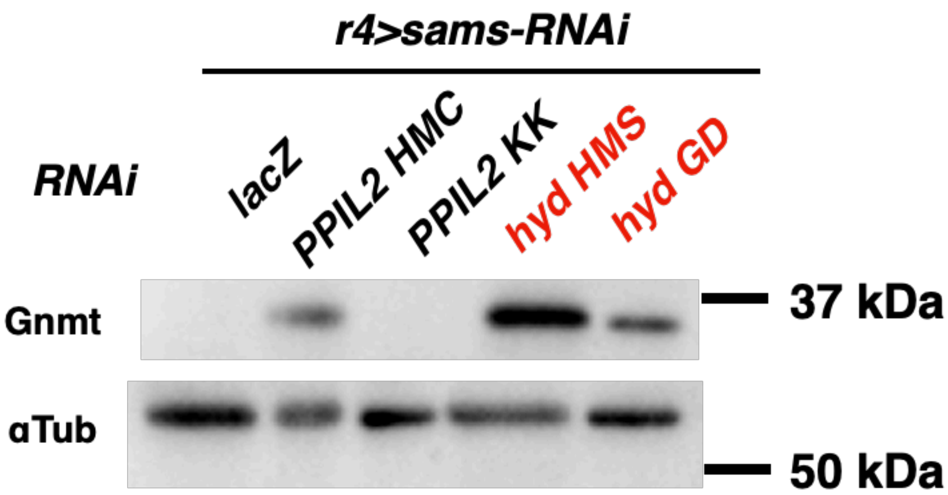


**Fig. S5. Confirmation of the role of E3 ligases in Gnmt rescue.**

Western blot analysis of Gnmt in the fat bodies of larvae with knockdown of *PPIL2* or *hyd*. αTub was used as a loading control.

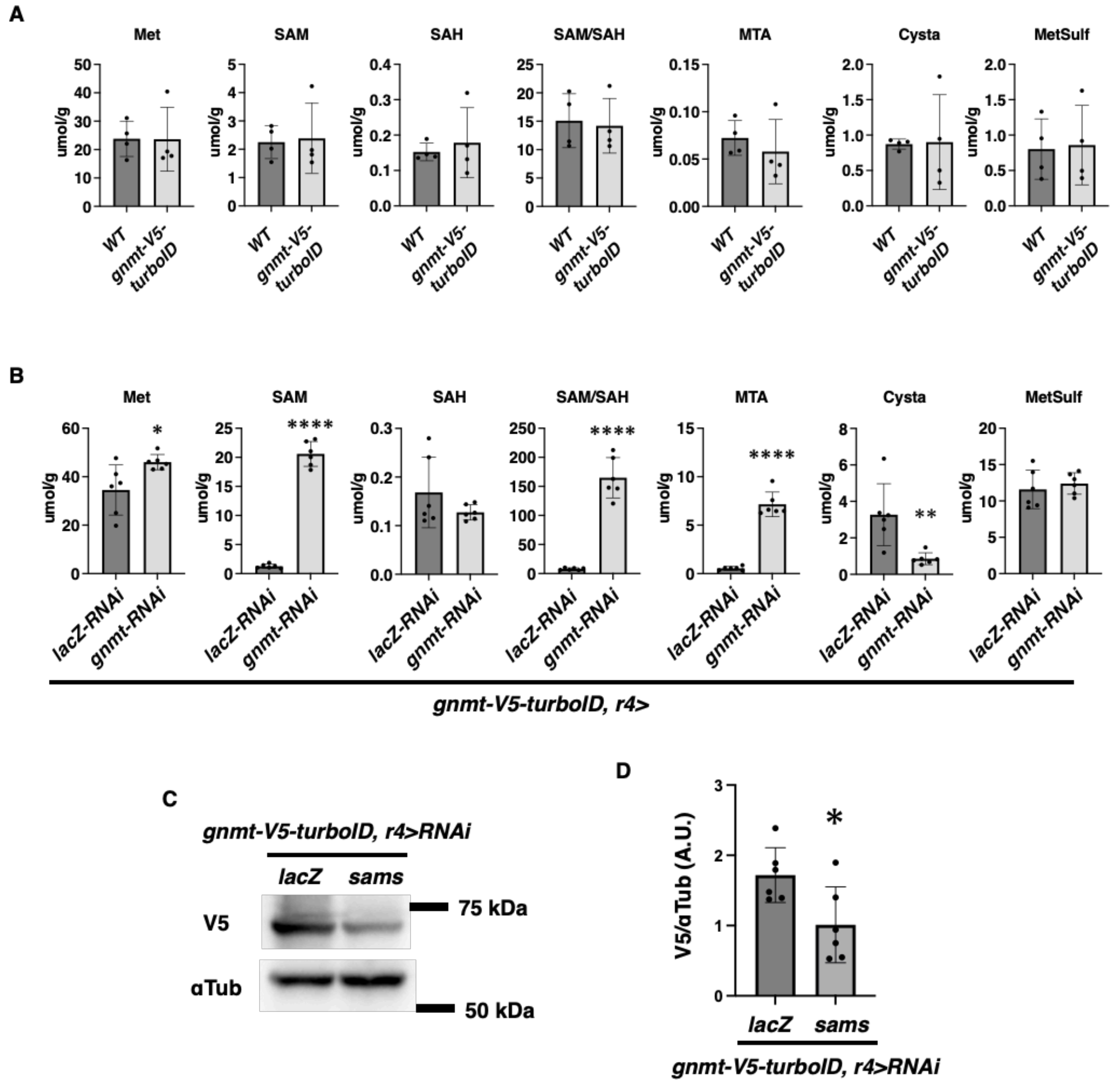


**Fig. S6. Verification of *gnmt-V5-turboID* knock in line.**

(A) The metabolite levels of SAM metabolism in $w^{iso31}$ and *gnmt-V5-turboID*. Metabolites were extracted from whole larvae. SEM was calculated from six independent samples. (B) The changes in metabolite levels of SAM metabolism caused by genetic manipulation of *gnmt* or *sams* in the fat body. Metabolites were extracted from whole larvae. SEM was calculated from six independent samples. Statistical significance was calculated using the two-tailed student's t-tests: *p < 0.05; **p < 0.01; ****p < 0.0001. (C) Western blot analysis of V5 in the fat body of larvae with knockdown of *sams*. αTub was used as a loading control. (D) Quantitative data on intensity of bands in (C). SEM was calculated from six independent samples. Statistical significance was calculated using the two-tailed student's t-tests: *p < 0.05.

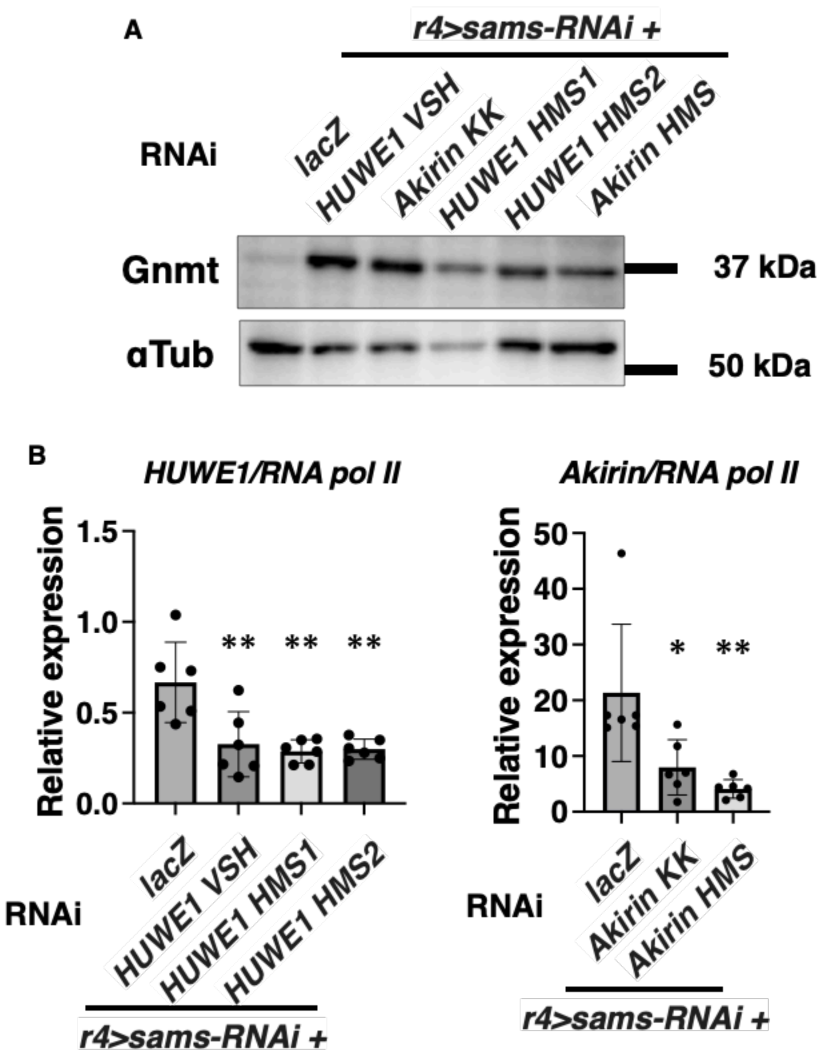


**Fig. S7. Confirmation of the contribution of *HUWE1* and *Akirin* in Gnmt rescue and the efficiency of knockdown.**

(A) Western blot analysis of Gnmt in the fat body of of larvae with genetic knockdown of *HUWE1* and *Akirin* in the fat body. αTub was used as a loading control. (B) qRT-PCR analysis of *HUWE1* and *Akirin* in the fat body of larvae with genetic knockdown of *HUWE1* and *Akirin* in the fat body. *RNA polII* was used as an internal control. SEM was calculated from six independent samples. One-way ANOVA Turkey's multiple comparison test was applied: $*p < 0.05$; $**p < 0.01$.

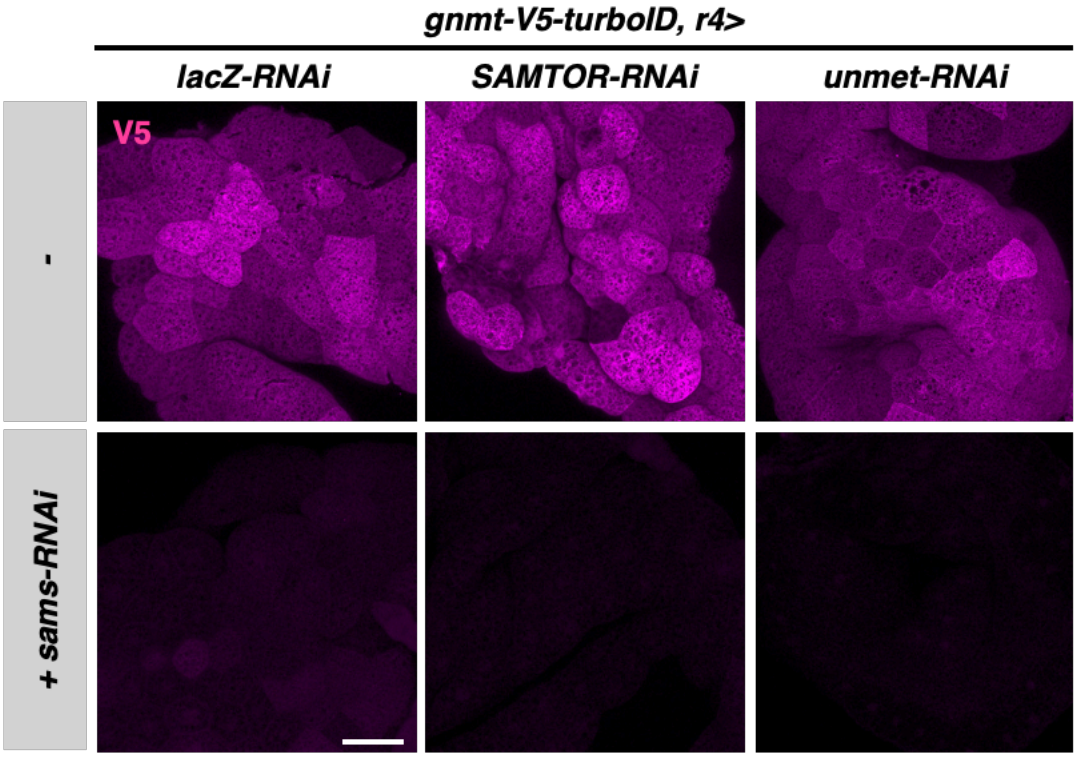


**Fig. S8. Knockdown of known SAM sensors does not rescue Gnmt decrease by *sams-RNAi*.**

Representative images of larval fat bodies expressing *gnmt-V5-turboID* with knockdown of *lacZ*, *SAMTOR*, or *unmet*. Fat bodies were stained with an anti-V5 antibody. (Scale bar, 75 µm.)

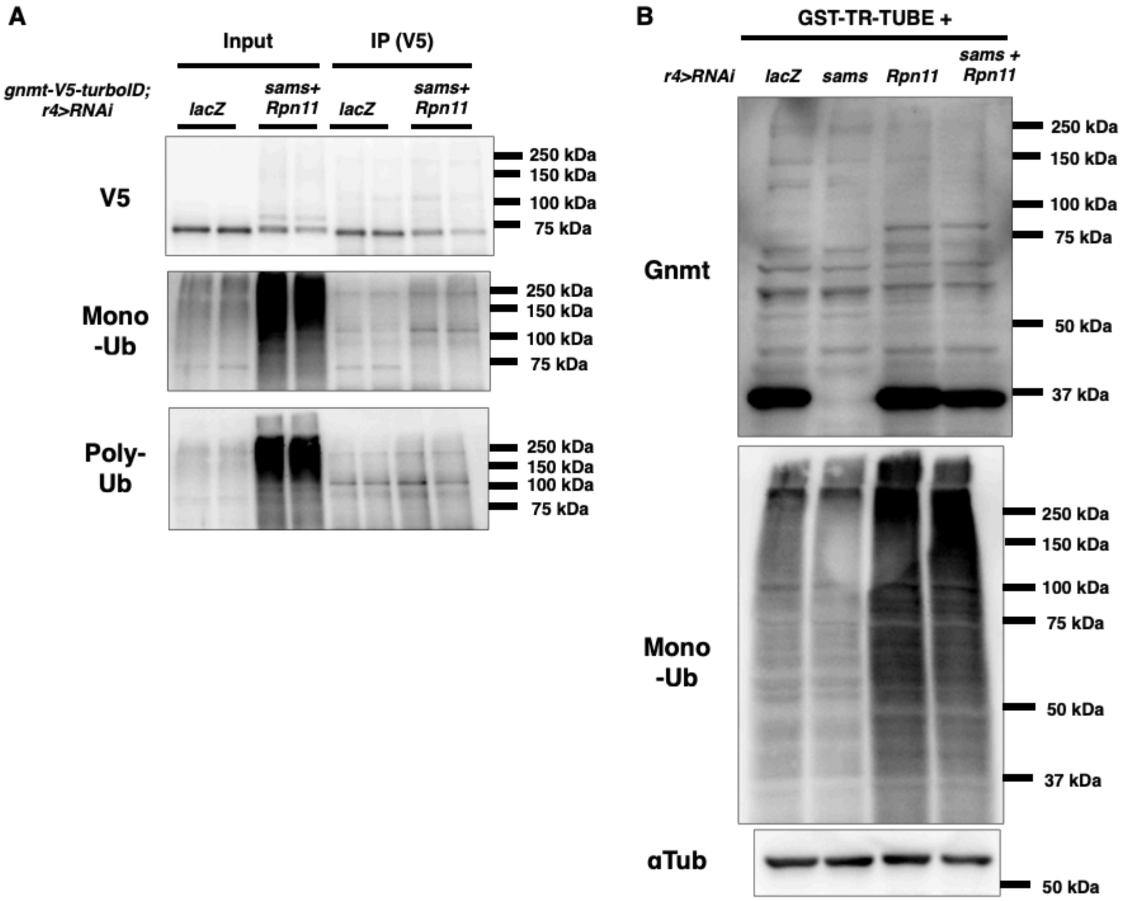


**Fig. S9. Ubiquitinated Gnmt is not detected.**

(A) Western blot analysis of V5, mono-ubiquitin, and poly-ubiquitin in the fat body of of *gnmt-V5-turboID* larvae with genetic knockdown of *sams* and *Rpn11* in the fat body. Immunoprecipitation (IP) was performed with V5 antibody. Duplicate samples are indicated. (B) Western blot analysis of Gnmt and mono-ubiquitin in the fat body of of larvae with genetic knockdown of *sams* and *Rpn11* in the fat body. GST-TR-TUBE was mixed in lysate samples for protecting linear ubiquitin chains. αTub was used as a loading control.

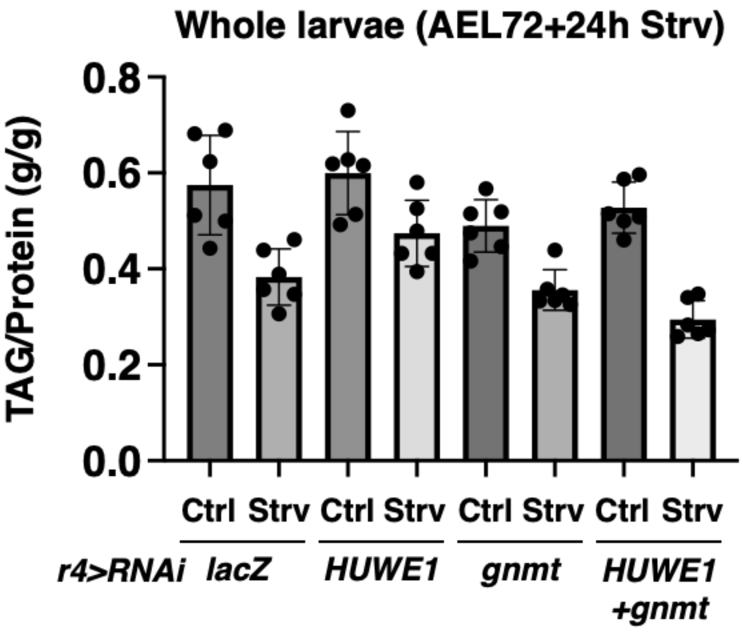


**Fig. S10. Gnmt is a major target of HUWE1 in the context of TAG level regulation.**

Quantitative data on TAG levels in larvae. SEM was calculated from six independent whole larvae samples.

### Table S1. Metabolite profiling of whole larvae under starvation (related to Fig. 1B)

| | Ctrl1 | Ctrl2 | Ctrl3 | Ctrl4 | Ctrl5 | Ctrl6 | Strv1 | Strv2 | Strv3 | Strv4 | Strv5 | Strv6 |
|---|---|---|---|---|---|---|---|---|---|---|---|---|
| 2-Ketoglutaric acid | 10.8022443 | 15.2542114 | 12.8332565 | 16.6457927 | 13.4975719 | 22.2247492 | 49.180488 | 31.8551096 | 57.1263079 | 38.450427 | 51.5665322 | 47.3695567 |
| 2-Morpholinoethanesulfonic acid | 3.28416827 | 2.53242884 | 3.26115736 | 1.89140378 | 3.28072353 | 1.50947254 | 2.52348034 | 1.09120982 | 1.84452834 | 3.63144991 | 4.52477907 | 3.79569461 |
| 3-HK | 41.0944305 | 35.6166768 | 38.5123316 | 32.518148 | 32.6514156 | 26.1255118 | 37.7308253 | 31.8494549 | 50.8609215 | 39.3023791 | 46.6730954 | 40.3288037 |
| 4-Hydroxyproline | 0.1245982 | 0.11936122 | 0.13876223 | 0.1136934 | 0.11542058 | 0.08717851 | 0.10746999 | 0.08514095 | 0.11956807 | 0.12327513 | 0.16068298 | 0.14628791 |
| 5-Glutamylcysteine | 6.17466534 | 9.22376287 | 6.82745799 | 7.8582641 | 6.95564131 | 6.93756636 | 4.27274383 | 5.73016139 | 8.17716261 | 9.87186299 | 7.33092516 | 21.7714361 |
| Acetylcarnitine | 4.67677001 | 4.10759932 | 4.87761482 | 3.83513726 | 4.14877839 | 3.2415298 | 3.0199027 | 2.99769611 | 4.19908087 | 3.02897173 | 3.83654182 | 3.23217496 |
| Acetylcholine | 0.15029002 | 0.09850896 | 0.16633649 | 0.03523633 | 0.03801814 | 0.01906747 | 0.04188965 | 0.02862525 | 0.05796061 | 0.04057106 | 0.03811134 | 0.0278996 |
| Adenine | 0.02130658 | 0.03620595 | 0.06361235 | 0.12730062 | 0.03458324 | 0.07061913 | 0.01419037 | 0.0381457 | 0.02557658 | 0.01161462 | 0.01960154 | 0.02450464 |
| Adenosine | 0.42636989 | 0.62222639 | 0.60307 | 0.4369305 | 0.67827642 | 0.4711568 | 0.28513019 | 0.40583319 | 0.51114256 | 0.24478805 | 0.29466817 | 0.3521617 |
| Alanine | 46.9849128 | 44.6378809 | 46.5330556 | 36.3447011 | 40.3667952 | 31.8050181 | 38.2252033 | 31.0568614 | 45.7468216 | 38.8415592 | 48.1512497 | 42.1756178 |
| Allantoic acid | 0.48066114 | 0.37758242 | 0.18464508 | 0.56857944 | 0.55977223 | 0.45411393 | 1.23051334 | 2.1564141 | 0.96865211 | 0.25059536 | 0.58119797 | 0.30428407 |
| Allantoin | 2.40368705 | 1.28133548 | 1.6057663 | 2.11504203 | 1.73860782 | 1.66911347 | 1.61802263 | 1.49033396 | 2.53417207 | 2.65710376 | 4.72411926 | 2.24311027 |
| AMP | 12.0013709 | 10.9784404 | 11.0977263 | 9.59897634 | 9.90692634 | 7.65883635 | 10.1475161 | 8.42080698 | 13.0749314 | 10.2764922 | 13.1094582 | 11.4721319 |
| Arginine | 12.8173604 | 11.807413 | 12.4855714 | 10.2702284 | 10.7667843 | 8.58561798 | 11.0157742 | 9.50045169 | 14.2876307 | 10.9767734 | 13.3204601 | 12.044226 |
| Argininosuccinic acid | 2.51169757 | 2.20928224 | 2.15261867 | 1.64178811 | 1.24484039 | 1.5491788 | 1.23067369 | 1.23455969 | 1.86587457 | 1.66113041 | 2.07562907 | 1.4797233 |
| Asparagine | 51.2197262 | 54.8699542 | 54.2741616 | 37.8468184 | 46.8518547 | 34.4721065 | 21.512718 | 16.4087849 | 25.4848232 | 18.0049379 | 25.6214914 | 18.5484885 |
| Aspartic acid | 11.1031104 | 8.18889012 | 9.45351645 | 7.08887506 | 8.57352007 | 4.68195453 | 4.32597775 | 2.14887442 | 3.30419239 | 3.88649086 | 3.0320352 | 2.94891352 |
| Biotin B7 | 0.03489131 | 0.07050856 | 0.08980567 | 0.03058813 | 0.04207757 | 0.05638652 | 0.03900347 | 0.04223502 | 0.06904704 | 0.05250607 | 0.05830837 | 0.05283949 |
| Carnitine | 10.4894619 | 10.8440216 | 12.0200608 | 9.84922926 | 9.59298393 | 7.7863557 | 14.5480133 | 14.1085097 | 21.7674183 | 14.1566551 | 16.5579889 | 14.4726662 |
| Carnosine | 89.7485493 | 72.1181577 | 78.2860408 | 67.8792306 | 59.7229079 | 49.429945 | 97.2183709 | 93.5895478 | 148.535782 | 103.18465 | 117.916593 | 103.585332 |
| Choline | 18.8650442 | 19.6863531 | 21.8025975 | 14.6682099 | 19.9814994 | 14.2123942 | 10.3098635 | 10.043341 | 16.2092292 | 7.96486318 | 12.5372422 | 8.34298089 |
| Citicoline | 0.16935631 | 0.21131419 | 0.21823312 | 0.20504546 | 0.23618867 | 0.175324 | 0.24468363 | 0.19561652 | 0.28401671 | 0.2848785 | 0.35828632 | 0.33610093 |
| Cysteine | 43.5710938 | 41.2302069 | 55.9268353 | 27.2392583 | 38.3165869 | 21.7176634 | 19.2459069 | 41.4747267 | 41.1932668 | 21.2807002 | 27.5439307 | 13.0787309 |
| Cystine | 0.00686386 | 0.0127736 | 0.01723948 | 0.0226787 | 0.01885294 | 0.01952073 | 0.03900347 | 0.05958269 | 0.05086141 | 0.04389524 | 0.0667941 | 0.03547297 |
| Cytidine | 0.15620057 | 0.644982 | 0.53050381 | 0.24646689 | 0.52315928 | 0.2851659 | 0.14927946 | 0.19711806 | 0.33074502 | 0.15367339 | 0.22271316 | 0.21327306 |
| Cytosine | 0.14123354 | 0.66824517 | 0.55745442 | 0.24706666 | 0.54415903 | 0.29196492 | 0.14282565 | 0.18957838 | 0.33434325 | 0.1447021 | 0.21705601 | 0.19886625 |
| DOPA | 1.56834526 | 2.58449605 | 4.19355935 | 7.89608693 | 5.9433205 | 8.73685834 | 9.30803954 | 2.23085253 | 5.52196372 | 7.9315012 | 6.76590466 | 13.4224487 |
| Dopamine | 0.05286129 | 0.07076234 | 0.0718089 | 0.04775647 | 0.02404433 | 0.03886016 | 0.15184495 | 0.09041234 | 0.06865804 | 0.2301296 | 0.14673859 | 0.85561669 |
| FAD | 0.82008875 | 1.10720007 | 1.03067148 | 0.76875178 | 1.07949661 | 0.72964034 | 0.84236269 | 0.78495799 | 1.30965694 | 0.79303798 | 0.93521662 | 0.87293969 |
| FMN | 0.14395048 | 0.15239492 | 0.14994339 | 0.12880004 | 0.16873965 | 0.12582715 | 0.16415129 | 0.14267897 | 0.23339843 | 0.13388849 | 0.1823191 | 0.14750661 |
| Formylkynurenine | 0.50468467 | 0.41937154 | 0.4324125 | 0.1270757 | 0.13880134 | 0.0533043 | 0.54360333 | 0.15695965 | 0.22265238 | 0.14506255 | 0.08843022 | 0.12874729 |
| Fumaric acid | 17.7590565 | 15.956886 | 18.0338332 | 14.0558099 | 16.6880511 | 11.4915181 | 12.0566417 | 10.9882298 | 17.737065 | 9.02415622 | 12.5556528 | 10.7400395 |
| GABA | 2.52823758 | 3.04273823 | 3.13032441 | 2.30880436 | 2.54257049 | 2.30272146 | 2.24564576 | 2.33445932 | 3.6865282 | 2.40947214 | 2.5173832 | 2.50743829 |
| Glutamic acid | 28.5658291 | 24.5170564 | 25.9643952 | 20.8565343 | 22.3995937 | 16.7273961 | 21.3107257 | 16.2305799 | 25.8355529 | 21.263118 | 26.0697463 | 22.5080551 |
| Glutamine | 18.4538321 | 17.6934183 | 19.1927985 | 14.8066437 | 16.7280209 | 12.8960138 | 14.2402346 | 11.2426303 | 18.7385388 | 15.2119831 | 18.3476836 | 15.684536 |
| Glutathione | 411.903721 | 371.713587 | 385.125766 | 262.878586 | 309.478633 | 213.872372 | 179.811042 | 214.106445 | 315.456212 | 188.726218 | 202.316097 | 112.291921 |
| Glycine | 39.7329545 | 50.684529 | 51.1406298 | 36.8989611 | 54.2674218 | 40.7794493 | 49.0651612 | 36.9926709 | 59.4311174 | 53.4343231 | 77.30072 | 60.0367141 |
| Guanosine | 2.43161916 | 2.82875935 | 3.3746172 | 2.2824146 | 2.93258822 | 2.1787376 | 1.87268762 | 1.29372706 | 2.1668613 | 1.28998335 | 1.23454945 | 1.29060668 |
| Guanosine monophosphate | 1.56920324 | 1.95161089 | 1.69793293 | 2.09967299 | 1.8612495 | 1.36243245 | 2.59703776 | 2.51010844 | 4.25879197 | 3.10867203 | 3.96293451 | 3.89867503 |
| Histamine | 0.1579642 | 0.65432957 | 0.54462504 | 0.22292602 | 0.53080975 | 0.27579837 | 0.14026016 | 0.1834444 | 0.30298034 | 0.13605121 | 0.18688453 | 0.19795222 |
| Histidine | 12.8808511 | 12.8463885 | 13.1710523 | 10.0323082 | 11.7338268 | 8.81757024 | 11.7329089 | 10.4010288 | 14.9164936 | 11.8454666 | 15.4586652 | 13.4339394 |
| Hypoxanthine | 6.87511312 | 10.084331 | 13.0530042 | 8.68121986 | 10.357172 | 10.2256921 | 8.77662229 | 9.92669953 | 11.2540347 | 6.85482628 | 9.16979744 | 8.42685378 |
| Inosine | 7.73109417 | 11.837782 | 14.2280062 | 10.3365027 | 11.7810177 | 10.2712304 | 10.5470511 | 10.7959039 | 13.4031479 | 8.10431867 | 10.8081386 | 10.051124 |
| Isoleucine | 25.0230266 | 47.2592173 | 50.1452837 | 29.7368769 | 38.6119496 | 32.376468 | 21.5370501 | 22.5074005 | 34.6588888 | 20.3431203 | 24.0545098 | 20.2492758 |
| Kynurenic acid | 1.64570673 | 1.28526907 | 1.71891432 | 2.70907409 | 2.91186169 | 2.29927663 | 1.16781918 | 1.20871391 | 2.21679883 | 1.72657276 | 1.75912628 | 1.4625744 |
| Kynurenine | 6.10888664 | 4.92565932 | 5.81620876 | 4.59309323 | 2.36063769 | 2.59338701 | 8.12695197 | 7.84603371 | 11.1878078 | 5.44389073 | 4.44061652 | 5.82252859 |
| L-Cystathionine | 1.29855727 | 1.76732428 | 1.33528019 | 0.83682538 | 1.92631751 | 1.24068471 | 0.38145633 | 0.33471733 | 0.39911131 | 0.31691882 | 0.376945 | 0.27829958 |
| Leucine | 39.4919089 | 75.9787021 | 83.0185678 | 51.3630626 | 68.5393303 | 57.0551489 | 33.1380767 | 33.7985276 | 53.8567363 | 31.6632458 | 35.1810369 | 31.8122881 |
| Lysine | 22.6158123 | 21.097201 | 22.8437374 | 17.984286 | 19.8393022 | 15.5172919 | 16.9468268 | 12.7978503 | 22.7845006 | 18.5148996 | 22.1502825 | 18.4602195 |
| Malic acid | 36.8593311 | 34.5027515 | 38.0474447 | 31.5236588 | 37.4016925 | 23.5414016 | 19.5382927 | 18.6291266 | 26.0896655 | 21.5894086 | 22.9836704 | 18.9309001 |
| Methionine | 24.5691536 | 36.6941 | 40.4400352 | 28.7408508 | 39.3569725 | 27.1151768 | 16.4998703 | 16.2314425 | 24.9245697 | 13.4496854 | 14.4893072 | 15.7013367 |
| Methionine sulfoxide | 5.03950616 | 7.06083779 | 11.5407855 | 2.68845709 | 12.2776345 | 5.25162154 | 12.6966914 | 9.84191001 | 16.5805758 | 13.7349644 | 13.4068061 | 16.2580229 |
| MTA | 1.95148234 | 2.06086321 | 2.07987429 | 1.40589212 | 2.4215682 | 1.36228136 | 2.19261227 | 1.53295235 | 2.25185722 | 1.36571865 | 1.60335609 | 1.85277705 |
| NAD | 7.50963922 | 6.76209637 | 7.31853818 | 5.06653458 | 5.34560819 | 4.02689186 | 1.76638012 | 3.93600365 | 6.21092671 | 5.05444063 | 2.45738761 | 5.57682932 |
| Niacin | 0.51865072 | 0.57523478 | 0.64080085 | 0.87828429 | 0.70657534 | 0.76019059 | 1.58499194 | 1.67384225 | 2.62602408 | 1.61423139 | 2.00943046 | 1.879545 |
| Nicotinamide | 25.5260906 | 22.4438002 | 22.8306852 | 20.5592746 | 20.293256 | 15.3293368 | 20.6328751 | 21.1030381 | 28.8283529 | 19.1313954 | 21.2798254 | 19.0861107 |
| Nicotinic acid | 0.45759093 | 0.53251514 | 0.56324546 | 0.84713388 | 0.62979743 | 0.6938624 | 1.46317124 | 1.52793655 | 2.36826469 | 1.52247592 | 2.00278083 | 1.80180914 |
| Octopamine | 0.0572942 | 0.08247852 | 0.07728811 | 0.04824379 | 0.02541048 | 0.04209347 | 0.14543123 | 0.09587541 | 0.07697286 | 0.24078301 | 0.15492658 | 0.87511594 |
| Ophthalmic acid | 0.31578541 | 0.29459634 | 0.474598 | 0.13089922 | 0.29868051 | 0.23433947 | 0.06962901 | 0.05364039 | 0.19148396 | 0.07096931 | 0.15408297 | 0.1067671 |
| Oxidized glutathione | 11.8434544 | 10.6368948 | 12.4257455 | 7.32814475 | 12.9883864 | 8.41189553 | 5.21961016 | 4.05072828 | 8.03478956 | 3.33111195 | 4.89120375 | 4.02145938 |
| Pantothenic acid | 3.163002 | 2.77364678 | 3.70029638 | 1.78490759 | 2.71209081 | 1.83150422 | 5.19315354 | 5.38883086 | 7.07717526 | 5.29938888 | 5.92993633 | 4.99485502 |
| Phenylalanine | 12.1031372 | 12.6252192 | 13.4088413 | 11.256021 | 12.4706917 | 9.33048812 | 9.5117555 | 8.0076258 | 12.2241455 | 9.41837231 | 10.7000572 | 9.82327001 |
| Proline | 44.0107147 | 43.9018341 | 47.0768343 | 34.0169665 | 36.0683643 | 30.3340129 | 28.2645271 | 23.7172645 | 37.884601 | 29.7190792 | 33.7721578 | 29.161217 |
| Pyridoxal-5'-phosphate | 4.99779865 | 4.43734421 | 3.66862385 | 3.03980829 | 3.70645645 | 2.60903986 | 2.96791144 | 2.09695921 | 4.14627448 | 2.87229457 | 3.16914671 | 2.62639243 |
| Riboflavin | 24.934559 | 23.5039664 | 24.0420817 | 36.8010865 | 37.5811662 | 29.1456051 | 30.5391148 | 29.8284995 | 46.7209683 | 38.1754409 | 43.8697282 | 38.2422968 |
| SAH | 0.18885159 | 0.1946916 | 0.22950337 | 0.15575208 | 0.24333171 | 0.16734648 | 0.11721084 | 0.13047491 | 0.21555317 | 0.11382324 | 0.17973865 | 0.13632066 |
| SAM | 1.37353545 | 1.89269163 | 2.14660489 | 1.63856436 | 2.4252373 | 1.56573818 | 1.60050514 | 1.57429284 | 2.4747041 | 1.51366483 | 1.68339984 | 2.04877064 |
| Serine | 28.1722055 | 29.149642 | 36.9433161 | 20.9241955 | 31.0772147 | 24.0379112 | 17.2645468 | 15.4403186 | 21.6889867 | 17.9252776 | 21.6190561 | 18.6176934 |
| Serotonin | 0.00438525 | 0.00304536 | 0.00320735 | 0.003861 | 0.00261521 | 0.00190372 | 0.00452969 | 0.00281141 | 0.00705058 | 0.00384484 | 0.00406918 | 0.00383021 |
| Succinic acid | 20.2650818 | 19.0154855 | 21.7437961 | 23.5433795 | 26.9247282 | 18.0485207 | 15.2072641 | 20.2480186 | 29.6288123 | 19.239892 | 21.6499719 | 19.8716954 |
| Taurine | 12.5653994 | 12.7926294 | 12.0958344 | 8.37676296 | 8.10305534 | 7.40866272 | 24.7642364 | 21.3132548 | 39.6990306 | 24.9599702 | 31.1526481 | 26.8257899 |
| Threonine | 44.755587 | 42.3441322 | 48.4331404 | 32.324648 | 40.2026614 | 29.655743 | 19.1315823 | 15.1139716 | 25.9622688 | 15.567871 | 20.5094106 | 16.8825212 |
| Tryptophan | 17.0971703 | 16.9473473 | 18.4821042 | 13.8496774 | 15.3501951 | 11.6718582 | 11.9781938 | 10.7786521 | 16.1814159 | 9.86877911 | 10.6624917 | 10.0832455 |
| Tyrosine | 98.0257496 | 89.0824236 | 94.9866429 | 90.623795 | 91.4438548 | 71.3045901 | 79.4183983 | 72.2473163 | 108.297698 | 87.27579 | 93.9918009 | 87.8308073 |
| Uracil | 6.63544987 | 12.387808 | 12.9844026 | 11.2089768 | 12.1713477 | 11.1160914 | 7.77295441 | 12.7969877 | 18.0246313 | 10.1959909 | 11.2483047 | 12.208011 |
| Uric acid | 2.44086631 | 2.20243018 | 3.03245583 | 11.6625133 | 9.55110152 | 8.30740218 | 14.4360537 | 13.821171 | 21.5370832 | 22.9576905 | 16.2055086 | 23.5430823 |
| Uridine | 6.80804745 | 13.3557251 | 13.71296 | 11.707121 | 12.6348645 | 12.0135013 | 8.51261731 | 13.6978205 | 19.2614485 | 10.9271109 | 11.5902647 | 12.511294 |
| Valine | 32.3767989 | 34.7919339 | 36.2704864 | 28.2184909 | 32.4765477 | 25.4819319 | 26.8958381 | 23.1786562 | 35.9073273 | 26.7311191 | 32.2467612 | 28.3321941 |
| Xanthine | 0.6544027 | 0.9846243 | 1.14760594 | 0.88653109 | 0.82465016 | 0.97745697 | 1.62668116 | 1.71141289 | 1.61390152 | 1.4457794 | 1.30824131 | 1.71371431 |

**Table S2. Fold changes of metabolites in whole larvae under starvation (related to Fig. 1B)**

| Metabolite | FoldChange (log2(Strv/Ctrl)) | -log10 (P Value) |
|---|---|---|
| Cystathionine | -2.009505473 | 2.945209896 |
| Ophthalmic acid | -1.435562332 | 2.082442667 |
| Adenine | -1.403950897 | 1.177367999 |
| Aspartic acid | -1.32115409 | 2.743299857 |
| Asparagine | -1.154405972 | 3.526603495 |
| Acetylcholine | -1.110273014 | 0.850405431 |
| Oxidized GSH | -1.106705055 | 3.105965853 |
| Threonine | -1.07077718 | 3.505526875 |
| Histamine | -1.056282154 | 1.252730289 |
| Cytosine | -0.997282472 | 1.195923122 |
| Methionine | -0.959002233 | 2.953797954 |
| Cytidine | -0.913692088 | 1.17856807 |
| Guanosine | -0.809066883 | 3.022656568 |
| Leucine | -0.774716503 | 2.033656831 |
| Choline | -0.739657103 | 2.681595466 |
| Glutathione | -0.688916297 | 1.85249028 |
| Malic acid | -0.660023178 | 2.838037581 |
| Isoleucine | -0.638487679 | 1.662635638 |
| Adenosine | -0.629045182 | 2.028322535 |
| Serine | -0.597475016 | 2.19536946 |
| NAD | -0.527129297 | 1.121262723 |
| Cysteine | -0.476956841 | 0.813050819 |
| Tryptophan | -0.425288961 | 1.752111657 |
| SAH | -0.401218791 | 1.282565241 |
| Kynurenic acid | -0.397692494 | 0.854623213 |
| Formylkynurenine | -0.382441304 | 0.261259665 |
| Proline | -0.367122362 | 1.55721737 |
| Fumaric acid | -0.362526262 | 1.267965317 |
| P5P | -0.329033935 | 0.903930991 |
| Acetylcarnitine | -0.29291674 | 1.420454233 |
| Lysine | -0.2783921 | 0.43329349 |
| Phenylalanine | -0.254389869 | 1.382835843 |
| Argininosuccinic acid | -0.2443145 | 0.596725507 |
| Valine | -0.12987796 | 0.525234772 |
| Hypoxanthine | -0.12362014 | 0.345785446 |
| Glutamine | -0.094181011 | 0.303308935 |
| MTA | -0.063083641 | 0.137477988 |
| Glutamic acid | -0.061617948 | 0.170074384 |
| Inosine | -0.055005402 | 0.141813427 |
| Biotin B7 | -0.046654729 | 0.059200739 |
| Succinic acid | -0.041753299 | 0.09701948 |
| SAM | -0.019339215 | 0.038969023 |
| Tyrosine | -0.017361534 | 0.060460284 |
| Alanine | -0.014548765 | 0.041407972 |
| GABA | -0.014124227 | 0.03422757 |
| FAD | 0.000605232 | 0.001236647 |
| Nicotinamide | 0.034566007 | 0.093534499 |
| 4-Hydroxyproline | 0.08692372 | 0.227082846 |
| Arginine | 0.092368949 | 0.336106007 |
| AMP | 0.118855424 | 0.40943351 |
| Uracil | 0.119492515 | 0.233712845 |
| Uridine | 0.123337205 | 0.238863236 |
| 2-Morpholinoethanesulfonic acid | 0.143802412 | 0.174596498 |
| Histidine | 0.16291771 | 0.63441811 |
| FMN | 0.20716108 | 0.670269945 |
| 3-HK | 0.256752533 | 1.068692757 |
| Glycine | 0.29801856 | 0.876640091 |
| Riboflavin | 0.369435475 | 1.309046088 |
| 5-Glutamylcysteine | 0.37810091 | 0.35992969 |
| Serotonin | 0.458676091 | 0.91604132 |
| Citicoline | 0.487067417 | 1.736118714 |
| Allantoin | 0.49756304 | 0.721238885 |
| DOPA | 0.547042228 | 0.60385678 |
| Carnitine | 0.658288501 | 2.472218813 |
| Carnosine | 0.670562317 | 2.545994276 |
| Kynurenine | 0.699473697 | 1.322175619 |
| Xanthine | 0.782755378 | 4.393808113 |
| MetSulf | 0.911858084 | 2.111057124 |
| GMP | 0.947888966 | 2.667426334 |
| Allantoic acid | 1.064729405 | 0.77988659 |
| Pantothenic acid | 1.085623072 | 4.335069553 |
| Taurine | 1.459632442 | 3.240430965 |
| Niacin | 1.480974737 | 3.471440793 |
| Nicotinic acid | 1.520799859 | 3.690076301 |
| Cystine | 1.593881727 | 3.286517742 |
| 2-Ketoglutaric acid | 1.594285672 | 3.802161734 |
| Uric acid | 1.596683329 | 3.311642448 |
| Octopamine | 2.255450005 | 0.815191427 |
| Dopamine | 2.334067977 | 0.819383322 |
